# Ergothioneine boosts mitochondrial respiration and exercise performance via direct activation of MPST

**DOI:** 10.1101/2024.04.10.588849

**Authors:** Hans-Georg Sprenger, Melanie J. Mittenbühler, Yizhi Sun, Jonathan G. Van Vranken, Sebastian Schindler, Abhilash Jayaraj, Sumeet A. Khetarpal, Ariana Vargas-Castillo, Anna M. Puszynska, Jessica B. Spinelli, Andrea Armani, Tenzin Kunchok, Birgitta Ryback, Hyuk-Soo Seo, Kijun Song, Luke Sebastian, Coby O’Young, Chelsea Braithwaite, Sirano Dhe-Paganon, Nils Burger, Evanna L. Mills, Steven P. Gygi, Haribabu Arthanari, Edward T. Chouchani, David M. Sabatini, Bruce M. Spiegelman

## Abstract

Ergothioneine (EGT) is a diet-derived, atypical amino acid that accumulates to high levels in human tissues. Reduced EGT levels have been linked to age-related disorders, including neurodegenerative and cardiovascular diseases, while EGT supplementation is protective in a broad range of disease and aging models in mice. Despite these promising data, the direct and physiologically relevant molecular target of EGT has remained elusive. Here we use a systematic approach to identify how mitochondria remodel their metabolome in response to exercise training. From this data, we find that EGT accumulates in muscle mitochondria upon exercise training. Proteome-wide thermal stability studies identify 3-mercaptopyruvate sulfurtransferase (MPST) as a direct molecular target of EGT; EGT binds to and activates MPST, thereby boosting mitochondrial respiration and exercise training performance in mice. Together, these data identify the first physiologically relevant EGT target and establish the EGT-MPST axis as a molecular mechanism for regulating mitochondrial function and exercise performance.

## Introduction

Exercise training can prevent and ameliorate many chronic diseases ^1,2^. Consistent with this, exercise capacity is a powerful predictor of mortality, with an increase in each one metabolic equivalent (1-MET) exercise capacity conferring a 12% improvement in survival ^3^. Thus, it is of great interest to identify key regulators of the beneficial effects of exercise. In recent years, defining the molecular signature of exercise adaptation using multi-omics approaches has provided a comprehensive overview of molecular changes in body fluids or tissues; several potentially important molecular mediators of the beneficial effects of exercise have been identified ^4–13^. Improved mitochondrial function is central to ensure metabolic adaptation to exercise training. Upregulation of mitochondrial biogenesis by the transcriptional co-activator PGC-1⍺, and regulation of mitochondrial dynamics and mitophagy ensure optimized functioning of the mitochondrial reticulum in response to exercise ^14,15^. However, our understanding of how mitochondria remodel their metabolome at the organellar level in response to training remains far from complete. One useful tool in this regard is the MITO-Tag Mouse, expressing a mitochondrially localized 3XHA epitope tag, which allows the study of mitochondrial metabolism *in vivo* ^16^. This Organelle-IP approach provides a higher resolution of compartmentalized metabolism compared to traditional methods and allows the analysis of the metabolome, due to the rapid isolation procedure ^17^.

Here, we isolated mitochondria from muscle tissue of MITO-Tag Mice after 4 weeks of endurance exercise training and analyzed their metabolome using mass spectrometry. We found muscle mitochondria are enriched in the metabolite ergothioneine (EGT) in response to endurance exercise training. The diet-derived, atypical amino acid EGT accumulates in human tissues at high levels but declines in human plasma with age and reduced EGT levels are linked to a multitude of age-related disorders, including neurodegenerative and cardiovascular diseases ^18–22^. Conversely, EGT supplementation is protective in animal models of several human diseases, improves acute aerobic performance and shows promising results in murine aging models ^22–26^. Together, this has led to the proposal that EGT be considered a “longevity vitamin” ^23^. However, its mechanism of action, and particularly its physiologically relevant, direct molecular target, has remained elusive ^22,27^. In this study, we have used a systematic proteome-wide solubility alteration assay and purified recombinant protein, to show that EGT binds the mitochondrial enzyme 3-mercaptopyruvate sulfurtransferase (MPST). We further demonstrate that EGT boosts mitochondrial respiration in mammalian cells via activation of MPST. Finally, enriching EGT in the diet augments endurance exercise training performance in an MPST-dependent manner.

## Results

### MITO-IP reveals accumulation of ergothioneine (EGT) in muscle mitochondria upon exercise training

To enable rapid isolation of mitochondria from muscle tissue we used MITO-tag Mice (Fig. 1A). MITO-tag Mice express the 3XHA-EGFP-OMP25 construct allowing epitope tagging of mitochondria; this was followed by HA-immunoprecipitation using magnetic beads (MITO-IP) ^16^. As expected, immunoblot analysis of MITO-IP from muscle tissue revealed substantial enrichment of mitochondria (Fig. 1B). Consistent with previous reports using MITO-Tag Mice, this rapid isolation workflow demonstrates minimal enrichment of endoplasmic reticular proteins and peroxisome marker proteins, organelles capable of forming contact sites with mitochondria (Fig. 1B) ^16^. To analyze how muscle mitochondria remodel their metabolism in response to exercise training we subjected MITO-Tag Mice to a 4 week voluntary wheel running (VWR) protocol before we performed MITO-IP from gastrocnemius muscle tissue followed by metabolomics (Fig. 1C). As expected, our running protocol, throughout which mice were running more than 200 km within 4 weeks, did not affect grip strength but resulted in a gene expression profile associated with muscle adaptation to endurance training (Fig. S1A-C). This includes increased expression of myosin heavy polypeptide 1 and 2 (*Myh1*, *Myh2*) (Fig. S1C) ^28^. Our exercise protocol allowed mice to train during their natural active phase, which is an important determinant of metabolic adaptations to exercise ^7^.

**Fig. 1.**
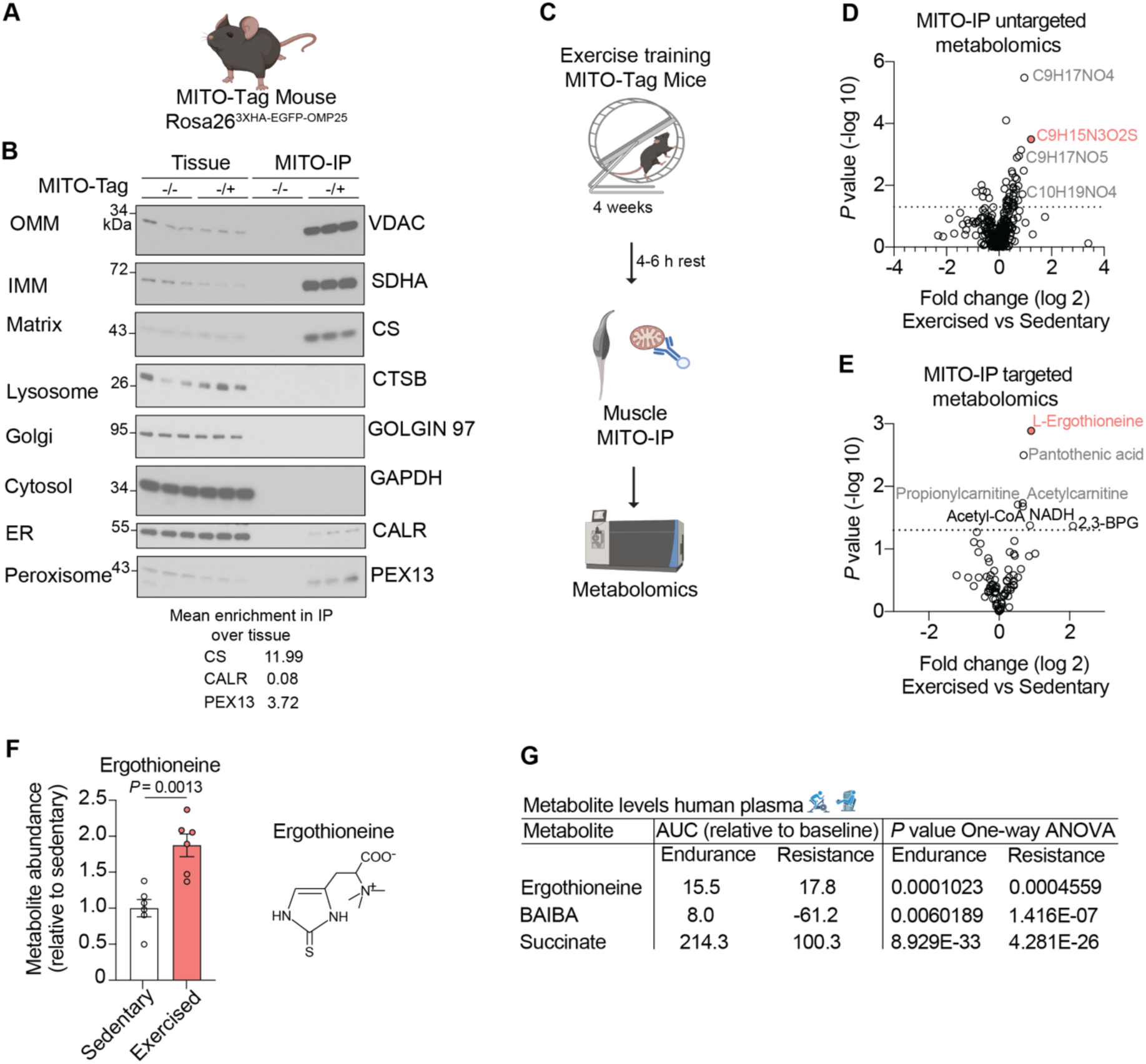
MITO-IP reveals accumulation of ergothioneine in muscle mitochondria upon exercise training. (**A**) Schematic of MITO-Tag Mouse expressing the 3XHA-EGFP-OMP25 construct knocked into the *Rosa26* locus. (**B**) Immunoblot analysis of gastrocnemius muscle tissue and the anti-HA immunoprecipitates (MITO-IP) from control mice (-/-, n = 3) and mice expressing one copy of the MITO-Tag (-/+, n = 3). OMM, outer mitochondrial membrane; IMM, inner mitochondrial membrane; Matrix, mitochondrial matrix; Golgi, Golgi complex; ER, endoplasmic reticulum. The mean enrichment in IP over tissue of the markers CS, CALR and PEX13 was quantified and is shown below. (**C**) Schematic of experimental set up for metabolomics from gastrocnemius muscle MITO-IP upon exercise training. (**D**) Volcano plot untargeted metabolomics of gastrocnemius muscle anti-HA immunoprecipitates (MITO-IP) in sedentary and exercised (4 weeks VWR) MITO-Tag Mice. Dotted line indicates metabolites are significantly different between exercised (-/+, n = 6) vs. sedentary (-/+, n = 6) mice. Labelled metabolites were subsequently validated using targeted metabolomics, see panel (E). (**E**) Volcano plot targeted metabolomics of gastrocnemius muscle anti-HA immunoprecipitates (MITO-IP) in sedentary and exercised (4 weeks VWR) MITO-Tag Mice. Metabolites included in this plot fulfilled criteria to be mitochondrial as described previously ^16^. Dotted line indicates metabolites are significantly different between exercised (-/+, n = 6) vs. sedentary (-/+, n = 6) mice. Ergothioneine is highlighted in red in (D) and (E). (**F**) Comparison of ergothioneine levels in sedentary (-/+, n = 6) vs exercised (-/+, n = 6) anti-HA immunoprecipitates (MITO-IP) from gastrocnemius muscle. Data are means ± s.e.m. Schematic of the metabolite ergothioneine (thione) is included in this panel. (**G**) Ergothioneine, BAIBA (β-aminoisobutyric acid) and succinate levels in human plasma as analyzed previously ^29^ upon endurance and resistance exercise. AUC = area under the curve. *P* values calculated using Compound Discoverer, Thermo Fisher Scientific (D), two-tailed unpaired t-test (E, F), one-way analysis of variance (ANOVA) (G).

Using polar metabolomics, we detected 81 metabolites in mitochondria from muscle tissue. These include known mitochondrial metabolites such as NAD, NADH, NADP, NADPH and ⍺-ketoglutarate and met criteria for being mitochondrial as defined previously (Fig. S1D) ^16^. Immunoblot analyses revealed that MITO-IPs from sedentary and exercised mice generally behaved similar with respect to mitochondrial enrichment (Fig. S1E). We performed a direct comparison of sedentary and exercised mice by untargeted and targeted, polar metabolomics, using our in-house library of over 100 metabolites. This showed that a distinct subset of metabolites belonging to fatty acid oxidation (FAO) pathways accumulates inside mitochondria with exercise training, i.e acetylcarnitine, propionylcarnitine, acetyl-CoA, pantothenic acid and NADH (Fig. 1D-E, Fig. S1D and F). Notably, the changes in those metabolites could only be observed in MITO-IPs from muscle and not when comparing metabolomics from muscle tissues without enriching for mitochondria (Fig. S1D). In our untargeted metabolomics dataset, we focused on the metabolite most highly enriched in mitochondria from exercised mice compared to sedentary mice, C9H15N3O2S. This metabolite could represent the diet-derived, atypical amino acid ergothioneine (EGT) (Fig. 1D). Using targeted metabolomics with a validated standard, we confirmed that EGT represents the most significantly enriched metabolite inside muscle mitochondria with exercise training (Fig. 1E). EGT accumulated almost two-fold in muscle mitochondria upon exercise training when compared to mitochondria from sedentary muscle tissues (Fig. 1F). Interestingly, analysis of publicly available untargeted human plasma metabolomics showed significant enrichment of EGT in human plasma upon endurance and resistance exercise (Fig. 1G) ^29^. Taken together, our data reveal exercise-dependent metabolic remodeling of mitochondria including substantial accumulation of the diet-derived, atypical amino acid EGT.

### PGC-1⍺ controls EGT levels in muscle

EGT is exclusively diet-derived and imported into certain gut microbes and mammalian cells through the EGT-specific transporters EgtUV and SLC22A4, respectively ^30,31^. How expression of *Slc22a4* is regulated remains poorly understood. Given the absence of mammalian EGT biosynthesis, we hypothesized that the increase of EGT in muscle mitochondria with training could be due to increased expression of the plasma membrane transporter SLC22A4. After an acute bout of exercise, the transcriptional co-activator PGC-1⍺ (*Ppargc1a*) is induced and controls expression of a broad set of nuclear genes encoding mitochondrial proteins, thereby inducing mitochondrial function and biogenesis ^32,33^. Thus, we speculated that SLC22A4 could be controlled by PGC-1⍺. Indeed, *Slc22a4* expression followed *Ppargc1a* expression in the gastrocnemius muscle upon an acute bout of exercise and during differentiation of C2C12 myoblasts into myotubes (Fig. 2A, Fig. S1G). Analysis of single-cell RNA sequencing data from human skeletal muscle tissue (Human Genotype Expression (GTEx) Project) ^34^ showed highest expression of both *PPARGC1A* and *SLC22A4* within the myocyte population (Fig. 2B). Furthermore, single-nucleus RNA sequencing revealed that expression of *PPARGC1A* and *SLC22A4* correlate in the human heart ^35^. To test if PGC-1⍺ expression is sufficient to induce *Slc22a4* expression, we used mice with forced expression of PGC-1⍺ in muscle (MCK-PGC-1⍺) (Fig. 2C) ^36^. Gene expression and immunoblot analysis of gastrocnemius muscle from MCK-PGC-1⍺ mice revealed a significant increase in *Slc22a4* mRNA and protein expression (Fig. 2D). In line with this, mass spectrometry showed a ∼28% increase in EGT metabolite levels in gastrocnemius muscle from MCK-PGC-1⍺ when compared to control animals (Fig. 2E), and EGT metabolite levels are depleted in hearts of cardiomyocyte-specific PGC-1⍺ knockout mice ^37^. Different isoforms of PGC-1⍺ have been described; PGC-1⍺1 is known to regulate oxidative metabolism in response to endurance exercise ^38^. We analyzed micro array data from primary myotubes overexpressing PGC-1⍺ isoforms 1-4 individually ^38^ and found that only PGC-1⍺1 expression induces *Slc22a4* expression, while PGC-1⍺2 and PGC-1⍺4 suppressed its expression (Fig. 2F). Our data are consistent with the transcriptional co-activator PGC-1⍺ contributing to the control of EGT level in muscle by regulating the expression of the EGT-specific transporter SLC22A4.

**Fig. 2.**
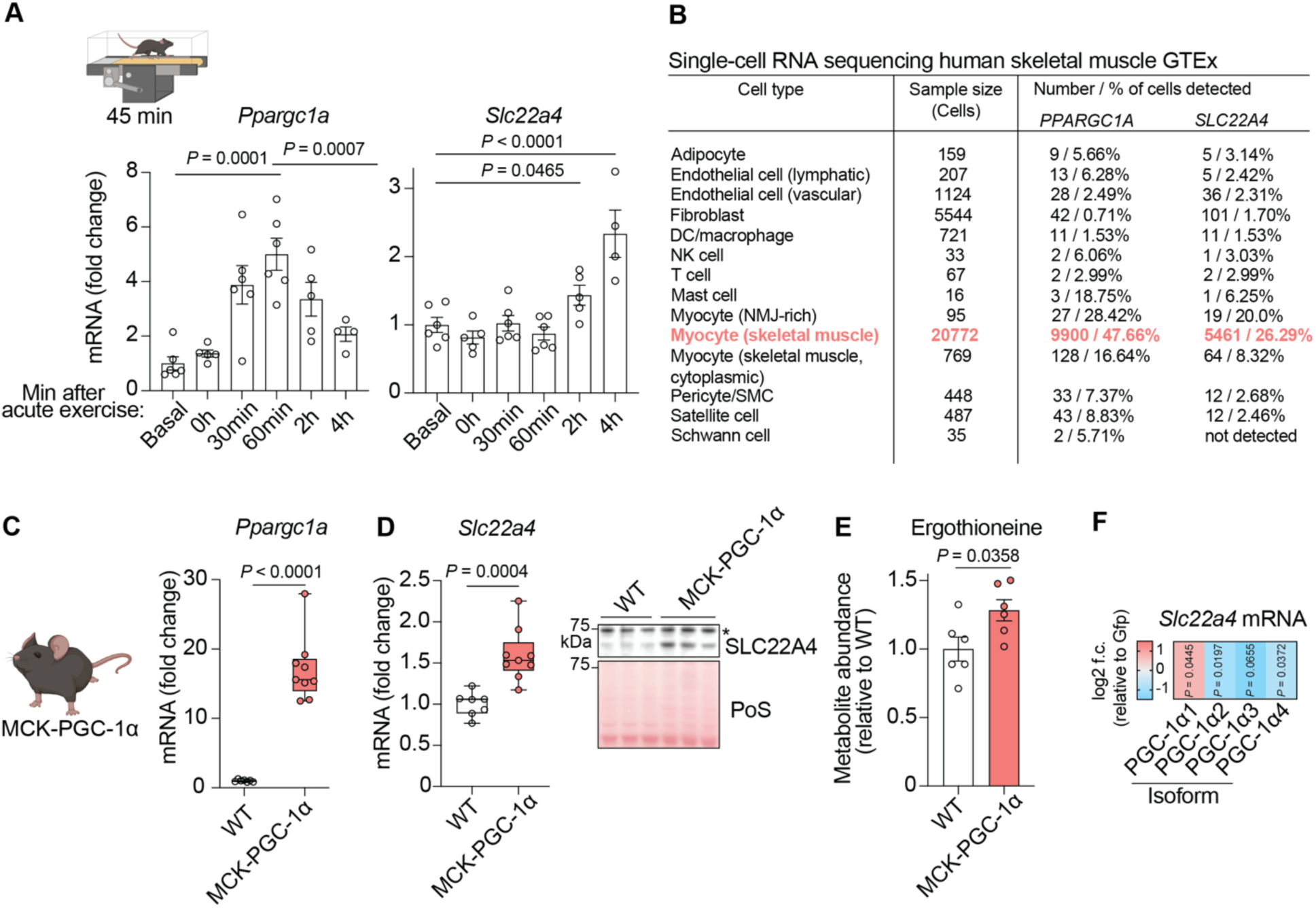
PGC-1⍺ controls ergothioneine levels in muscle. (**A**) *Ppargc1a* and *Slc22a4* expression in gastrocnemius muscle at different timepoints after an acute bout of exercise monitored by RT-qPCR (n = 4-6 per timepoint). Data are means ± s.e.m. (**B**) Analysis of single-cell RNA sequencing data from human skeletal muscle. (The Genotype-Tissue Expression project, GTEx ^34^). Myocytes are highlighted in red. (**C**) Schematic of MCK-PGC-1⍺ Mice with forced expression of PGC-1⍺ in muscle tissue and *Ppargc1a* expression in gastrocnemius muscle from WT (n = 7) and MCK-PGC-1⍺ Mice (n = 9) monitored by RT-qPCR. (**D**) *Slc22a4* expression in gastrocnemius muscle from WT (n = 7) and MCK-PGC-1⍺ Mice (n = 9) monitored by RT-qPCR (left panel). Immunoblot analysis of SLC22A4 protein levels in gastrocnemius muscle from WT (n = 3) and MCK-PGC-1⍺ Mice (n = 3) (right panel). * = indicates unspecific binding. Center lines denote medians; box limits denote 25^th^ and 75^th^ percentiles; whiskers denote maxima and minima in (C, D). (**E**) Ergothioneine level in gastrocnemius muscle from WT (n = 6) and MCK-PGC-1⍺ Mice (n = 6). Data are means ± s.e.m. (**F**) Heat map of changes in *Slc22a4* expression in myotubes expressing different PGC-1⍺ isoforms as analyzed previously ^38^. The data are presented as the log_2_-transformed fold change (f.c., relative to GFP expressing myotubes). *P* values calculated using one-way analysis of variance (ANOVA) (A), two-tailed unpaired t-test (C, D, E), NCBI GEO2R (F).

### EGT directly targets 3-mercaptopyruvate sulfurtransferase (MPST) to increase mitochondrial respiration

Exercise training induces mitochondrial biogenesis and turnover of damaged mitochondria via mitophagy ^15^. It has been noted, that the improvement of mitochondrial function (mainly increased respiratory capacity during β-oxidation and maximal oxidative phosphorylation, OXPHOS) observed with exercise training cannot solely be explained by an increased mitochondrial mass ^15^. To address if EGT could contribute to enhanced mitochondrial respiration and OXPHOS, we treated cultured cells with EGT and measured oxygen consumption rates (OCRs). A wide range of EGT concentrations has been reported in different body fluids and tissues, from ∼810 nM in human plasma to ∼0.86 ng per µl, ∼6.57 or ∼64.88 ng per mg in mouse plasma, heart or erythrocytes, respectively ^22^. We quantified EGT in mouse gastrocnemius muscle by mass spectrometry and determined an average EGT concentration of 8.89 ng per mg tissue in sedentary mice (Fig. S2A). EGT is known to protect cultured cells against superoxide-mediated cell death in a dose-dependent manner between 100 µM and 1 mM ^39^. Therefore, we initially decided to treat cells with 500 µM EGT. EGT was efficiently taken up by cells as determined by mass spectrometry and was sufficient to boost basal (∼30% in C2C12 myotubes, ∼23% in primary inguinal white adipocytes (primary iWAT), ∼67% in HeLa cells) and maximal (∼22% in C2C12 myotubes, ∼19% in primary iWAT, ∼68% in HeLa cells) mitochondrial respiration in a variety of different cell types (Fig. 3A, Fig. S2B-E). Immunoblot analysis using an OXPHOS antibody cocktail showed that the boost in respiration was independent of changes in expression of these mitochondrial respiratory chain subunits (Fig. S3A).

**Fig. 3.**
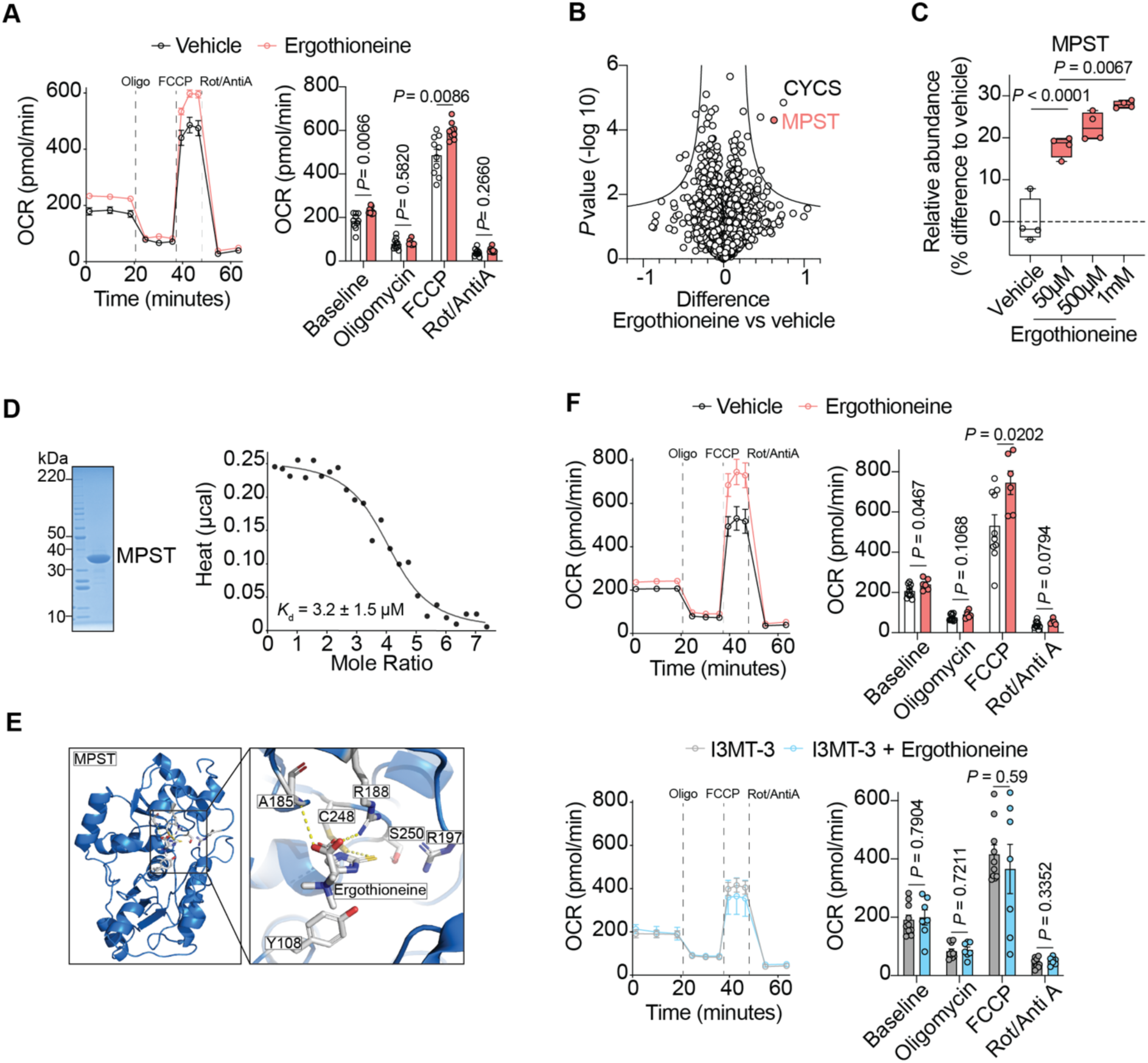
Ergothioneine directly targets 3-mercaptopyruvate sulfurtransferase (MPST) to increase mitochondrial respiration. (**A**) Oxygen consumption rates (OCR) of C2C12 myotubes after treatment with oligomycin (Oligo), carbonyl cyanide p-triflouromethoxyphenylhydrazone (FCCP), rotenone (Rot), antimycin A (AntiA). Equal amounts of cells were treated with vehicle (n = 10) or 500 µM ergothioneine (n = 9) for 72 h. (**B**) Volcano plot of proteins detected in proteome integral solubility alteration (PISA) assay (1 mM ergothioneine vs vehicle). Curved line indicates proteins significantly different between groups according to permutation-based FDR. CYCS = cytochrome C, MPST = 3-mercaptopyruvate sulfurtransferase. (**C**) Quantification for MPST detected in PISA assay across different ergothioneine concentrations (vehicle, 50 µM, 500 µM and 1 mM ergothioneine, n = 4). Center lines denote medians; box limits denote 25^th^ and 75^th^ percentiles; whiskers denote maxima and minima (**D**) SDS-gel of recombinant human MPST (left panel). ITC profile for the binding of ergothioneine to recombinant human MPST. n = 4.1 ± 0.2, Δ*H* = 0.204 ± 0.011 kcal *mol^-1^, Δ*S* = 25.83 kcal*mol^-1^*K^-1^, confidence level = 95% (right panel). Data are representative of two independent experiments. (**E**) Molecular modeling of the favorable binding mode of ergothioneine in the active site of MPST (PDB: 4JGT). (**F**) OCR of C2C12 myotubes after treatment with Oligo, FCCP, Rot, AntiA. Equal amounts of cells were treated with vehicle (n = 10) or 10 µM ergothioneine (n = 6) for 72 h. For I3MT-3 treatment cells were pre-treated with I3MT-3 for 4 h before the measurement (I3MT-3, n = 10; I3MT-3 + ergothioneine, n = 7). Data are means ± s.e.m (A, F). *P* values calculated using two-tailed unpaired t-test (A right panel, F right panels), one-way analysis of variance (ANOVA) (C).

Based on its chemical properties a variety of potential biological activities have been proposed for EGT in living cells, including a role as a protein ligand ^27^. We therefore used proteome integral solubility alteration (PISA) ^40,41^ to identify protein-ligand interactions, which could explain how EGT controls mitochondrial respiration. This method depends upon ligand binding affecting the thermostability of any protein in a given cellular proteome. Strikingly, the two proteins most stabilized in our PISA experiments using whole cell lysates are both known regulators of mitochondrial respiration, 3-mercaptopyruvate sulfurtransferase (MPST) and cytochrome c (CYCS) (Fig. 3B and C, Fig. S3B). While the thermostability of both proteins changed significantly upon treatment with increasing concentrations of EGT (500 µM, 1 mM) in the PISA assay, we did not detect any differences in absolute protein amounts of MPST and CYCS in whole cell lysates (Fig. 3C, Fig. S3B and C). Furthermore, in contrast to CYCS, MPST was stabilized at the lowest dose EGT (50 µM) in the PISA assay, suggesting a more effective engagement (Fig. 3C, Fig. S3B). A high degree of structural similarity between MPST and the bacterial ergothioneine synthase Eanb further supports binding of EGT to MPST (RMSD: 1.68 Å) (Fig. S3D). To experimentally test if EGT directly targets MPST, we purified recombinant human MPST protein (mitochondrial isoform MPST2) expressed in bacteria and measured binding to EGT by isothermal titration calorimetry (ITC) (Fig. 3D). Our experiments demonstrate that EGT binds recombinant MPST with a K_d_ of 3.2 µM in an entropy-dominated process (Fig. 3D). Protein-detected nuclear magnetic resonance spectroscopy (NMR) using ^15^N-labelled recombinant human MPST further supported direct binding of EGT to MPST (Fig. S3E-G). We acquired ^1^H^15^N-HSQC spectra of MPST in the presence and absence of its substrate 3-mercaptopyruvate (3-MP), known to bind to the active site of MPST. Here, we used chemical shift perturbation (CSP) and line broadening of the protein resonances as an indication of ligand binding to the protein. When EGT was added to MPST, we observed line broadening and signal enhancement for the same resonances of MPST. We further validated our findings with glutathione and L-cysteine, two known small molecule acceptors of MPST *in vitro* ^42^, that both elicited similar effects on the chemical environment of MPST, which translated to similar perturbation in the ^1^H^15^N-HSQC spectra of MPST (Fig. S3F). To further enhance the spectral resolution and reaffirm our findings, ^1^H^15^N-HSQC spectra on 600 µM MPST samples, with and without a twofold molar excess of EGT, were recorded on an 800 MHz spectrometer, corroborating the reproducibility of the observed spectral effects observed on the 600 MHz spectrometer (Fig. S3G).

MPST is one of three mammalian hydrogen sulfide (H_2_S)-producing enzymes ^42,43^. MPST uses 3-MP as a substrate to produce pyruvate and hydrogen sulfide (H_2_S) inside mitochondria and thereby can contribute to electron flow and mitochondrial respiration ^44,45^. Upon desulfuration of 3-MP, pyruvate and an enzyme bound persulfide is generated at the active site cysteine C248. Subsequently, pyruvate leaves the active site of MPST, allowing an acceptor molecule to enter. Upon transfer of the outer sulfur atom from the persulfide to the acceptor molecule, the acceptor molecule is released from the active site and releases H_2_S ^46^. We hypothesized that EGT can act as such an acceptor molecule on MPST and thereby boosts mitochondrial respiration. To test this, we first explored the molecular basis for direct action of EGT on the active site of human MPST using structural modeling. This method allowed us to predict the energetically favored binding sites and orientations of EGT to MPST. Strikingly, the predicted poses of EGT between nine variants of AutoDock were found to be in complete agreement with each other and in structural agreement with results obtained from Glide docking (Fig. 3E, Fig. S3H and I); directly interacting residues: R188, A185 and Y108. This analysis placed EGT in close proximity to the active site cysteine persulfide (248Cys-SSH) (Fig. 3E, Fig. S3H and I).

Based on our EGT quantification in mouse muscle tissue and our ITC data, we next treated C2C12 myotubes with a physiologically relevant dose of 10 µM EGT in combination with the MPST-specific inhibitor I3MT-3 and measured OCRs. I3MT-3 targets the active site cysteine persulfide of MPST and thereby prevents action of an acceptor molecule ^47^. While EGT treatment increased mitochondrial respiration in control cells, EGT was unable to boost mitochondrial respiration if cells were also treated with I3MT-3 (Fig. 3F). Importantly, similar results were obtained when inhibiting MPST in primary iWAT cells and upon deletion of *MPST* using CRISPR-Cas9 in human HeLa cells, strongly suggesting that EGT boosts mitochondrial respiration by activating MPST (Fig. S2D - F). Thus, using a combination of a systematic proteome-wide solubility alteration assay, pharmacological, genetic and biophysical experiments, we demonstrate that the diet-derived atypical amino acid EGT binds mitochondrial MPST and uses this enzyme to boost mitochondrial respiration.

### EGT-enriched diet augments exercise training performance via MPST in mice

The observation that EGT accumulates inside muscle mitochondria with exercise training and boosts mitochondrial respiration in different cell types via MPST, raised the intriguing possibility that increasing the availability of EGT during exercise training could further enhance endurance exercise performance in an MPST-dependent manner. Humans and mice obtain EGT exclusively from their diet. Despite only trace levels of EGT in standard mouse diets (5.6 ng/mg), rodents can show high basal level of EGT in their tissues (mouse liver 80.7 ng/mg), indicating efficient EGT uptake ^22,48^. To test if increasing the availability of EGT before and during exercise training could further enhance exercise performance, we generated a mouse diet enriched in EGT (209 ng/mg) and gave mice ad libitum access to either a control or EGT-enriched diet during exercise training (Fig. S4A). Importantly, neither body weight nor food consumption was affected when feeding the EGT-enriched diet for two weeks during a sedentary period prior to the exercise training protocol (Fig. 4A). However, the average running distance per hour of mice fed the EGT-enriched diet was significantly increased when individually housed mice had access to running wheels. On average control diet-fed mice ran 0.73 km per hour during the dark phase of an entire week, while EGT diet-fed mice ran 0.87 km per hour, reflecting a ∼19% increase in exercise performance during their active phase (Fig. 4B). Moreover, at peak activity EGT diet-fed mice were able to run ∼28% faster (Average 24.8 m per min) than control diet-fed mice (Average 19.4 m per min) (Fig. 4C, Fig. S4B).

**Fig. 4.**
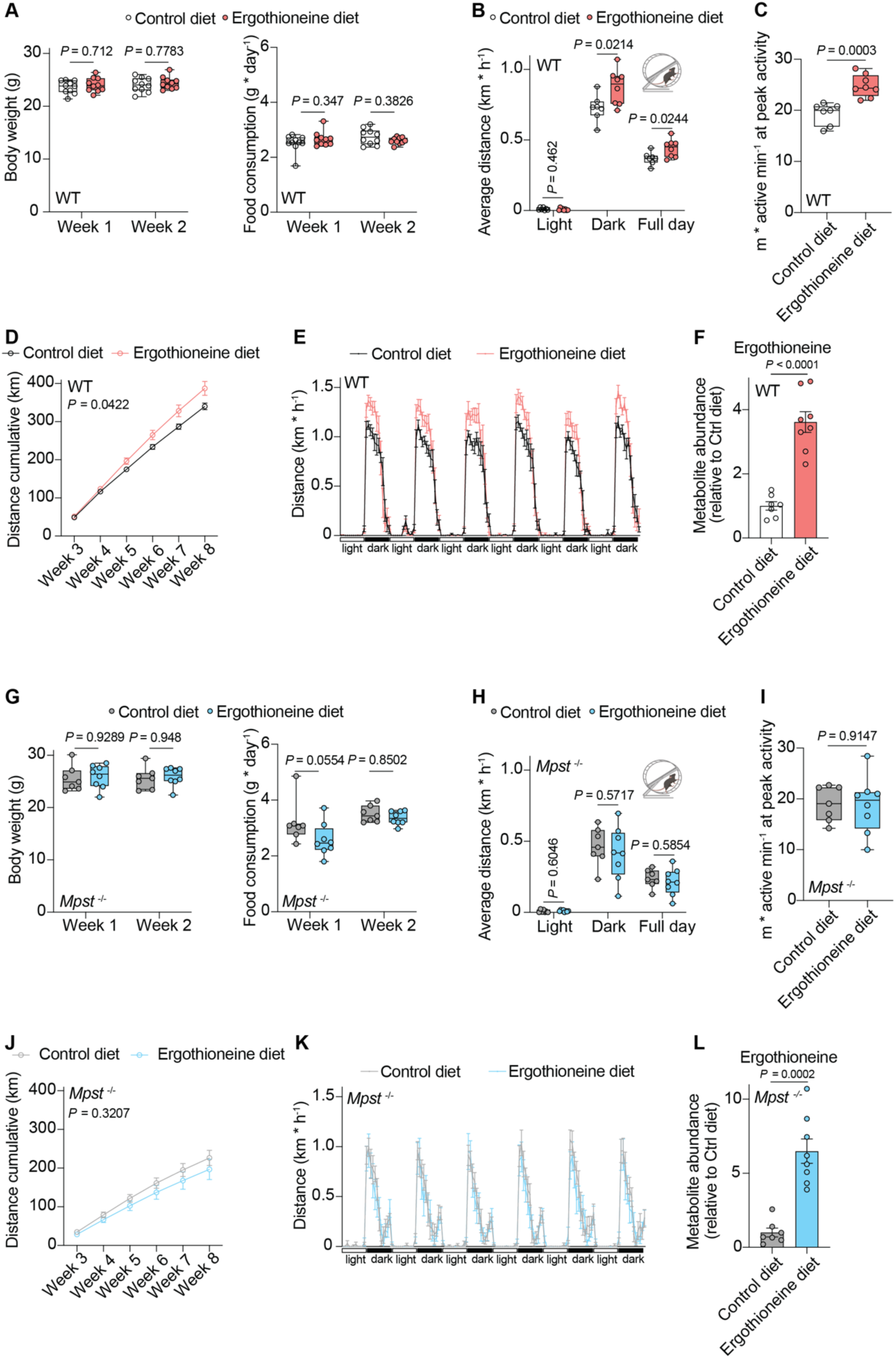
Ergothioneine-enriched diet augments exercise training performance via MPST in mice. (**A**) Body weight (left panel) and food consumption (right panel) of control-fed (n = 10) and ergothioneine-fed (n = 10) mice during the sedentary time period of 2 weeks. (**B**) Average running distance (km/h) of control-fed (n = 7) and ergothioneine-fed (n = 8) mice during 6 consecutive light and dark cycles. (**C**) Running speed at peak activity (m/active min) of control-fed (n = 7) and ergothioneine-fed (n = 8) mice. Data are average values from 3 timepoints. (**D**) Cumulative running distance (km) during weeks 3 - 8 of control-fed (n = 7) and ergothioneine-fed (n = 8) mice. (**E**) Running distance (km/h) of control-fed (n = 7) and ergothioneine-fed (n = 8) mice during 6 consecutive light and dark cycles. (**F**) Ergothioneine levels in gastrocnemius muscle of control-fed (n = 7) and ergothioneine-fed (n = 8) mice after week 8 of the VWR exercise protocol. (**G**) Body weight (left panel) and food consumption (right panel) of *Mpst*^-/-^ control-fed (n = 7) and ergothioneine-fed (n = 8) mice during the sedentary time period of 2 weeks. (**H**) Average running distance (km/h) of *Mpst*^-/-^ control-fed (n = 7) and ergothioneine-fed (n = 8) mice during 6 consecutive light and dark cycles. (**I**) Running speed at peak activity (m/active min) of *Mpst*^-/-^ control-fed (n = 7) and ergothioneine-fed (n = 8) mice. Data are average values from 3 timepoints. (**J**) Cumulative running distance (km) during weeks 3 - 8 of *Mpst*^-/-^ control-fed (n = 7) and ergothioneine-fed (n = 8) mice. (**K**) Running distance (km/h) of *Mpst*^-/-^ control-fed (n = 7) and ergothioneine-fed (n = 8) mice during 6 consecutive light and dark cycles. (**L**) Ergothioneine levels in gastrocnemius muscle of *Mpst*^-/-^ control-fed (n = 7) and ergothioneine-fed (n = 8) mice after week 8 of the VWR exercise protocol. Center lines denote medians; box limits denote 25^th^ and 75^th^ percentiles; whiskers denote maxima and minima (A, B, C, G, H, I). Data are means ± s.e.m. (D, E, F, J, K, L). *P* values calculated using two-way analysis of variance (ANOVA) (A, D, G, J), two-tailed unpaired t-test (B, C, F, H, I, L).

Finally, the cumulative running distance of mice fed the EGT-enriched diet was significantly increased (∼14%) over an entire 6 weeks of voluntary wheel running (VWR) (Fig. 4D, Fig. S4C). While the average total running distance of control diet-fed mice was 340 km, EGT diet-fed mice ran 387 km on average (Fig. 4D, Fig. S4C). Calculating the average running distance per hour during an entire week showed that exercise training performance was restricted to the dark phase in both groups, excluding significant disruption of the circadian running behavior upon EGT feeding (Fig. 4E). Feeding the EGT-enriched diet did not affect body weight or grip strength during the endurance exercise protocol and resulted in a ∼3.6-fold increase of EGT in gastrocnemius muscle tissue (Fig. S4D and Fig. 4F). Strikingly, when we subjected *Mpst*^-/-^ mice to the same diet and exercise protocol, we did not observe any difference between the control diet-fed and EGT diet-fed groups when analyzing their exercise performance (Fig. 4G-L, Fig. S4E-H). These data demonstrate that increasing the availability of EGT before and during exercise training improves endurance exercise performance via MPST in mice.

## Discussion

In this study, we identify the first physiologically relevant molecular target of ergothioneine (EGT), 3-mercaptopyruvate sulfurtransferase (MPST). This data also establishes a previously unidentified EGT-MPST axis as a key molecular mechanism regulating mitochondrial function and improvement in exercise performance upon training (Fig. 5). EGT is a sulfur-containing atypical amino acid and accumulates to high levels in human tissues but its synthesis has not been observed in mammals. EGT is believed to be exclusively diet-derived, mainly acquired from mushrooms and imported into tissues through its plasma membrane transporter SLC22A4 ^22,30,49^. Interestingly, EGT levels in human plasma decrease with advanced age and low levels of EGT in human plasma are associated with a range of age-related disorders ^18,22^. Conversely, EGT supplementation shows promising results in animal models mimicking a broad spectrum of human diseases including Alzheimer’s disease and depression and during aging in mice^22,26,50,51^.

**Fig. 5.**
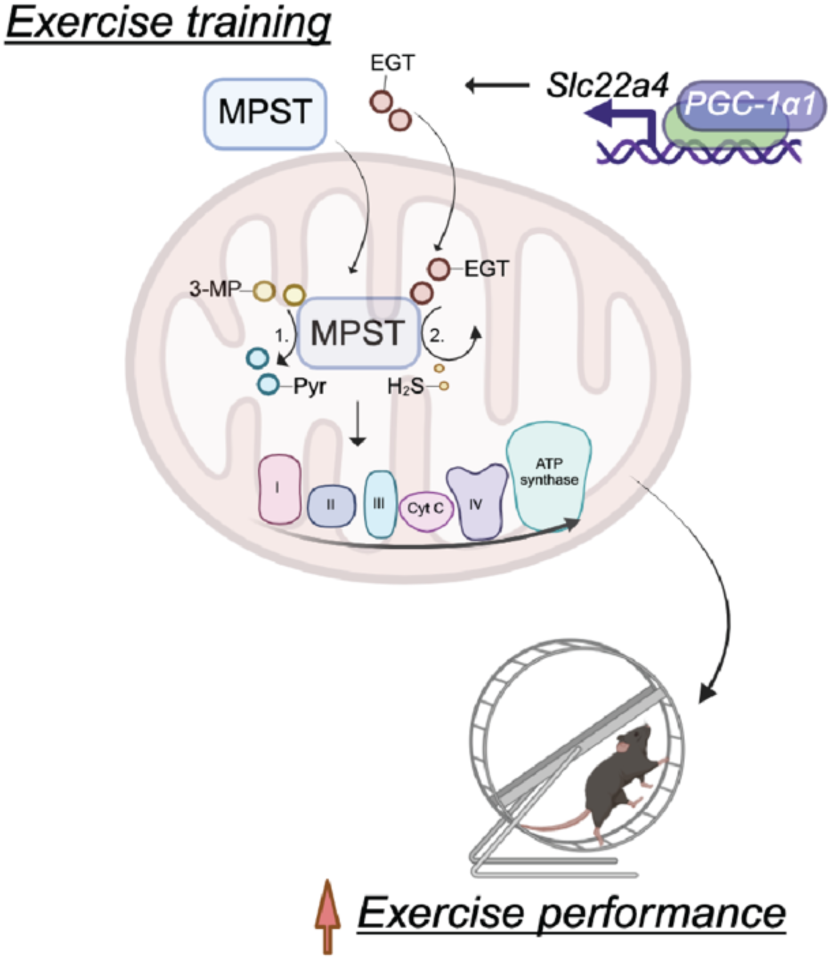
Ergothioneine boosts mitochondrial respiration and exercise performance via direct activation of MPST. Proposed model showing that upon exercise training PGC-1⍺ controls expression of the ergothioneine transporter SLC22A4, which in turn controls ergothioneine (EGT) levels within the tissue. Both, EGT and 3-mercaptopyruvate sulfurtransferase (MPST) accumulate inside mitochondria with exercise training. In mitochondria EGT binds to and activates MPST. MPST produces pyruvate (Pyr) and hydrogen sulfide (H_2_S) from 3-mercaptopyruvate (3-MP) and boosts mitochondrial respiration. Finally, increasing EGT improves endurance exercise training performance via MPST in mice.

Exercise training increases levels of fatty acid oxidation intermediates and EGT inside mitochondria, and boosts respiration of individual mitochondria. For instance, mean electron transport system capacity increases by ∼70% when comparing active individuals to elite athletes. Mean maximal exercise capacity increases by ∼25% from active to well-trained individuals and up to ∼51% when comparing active individuals to elite athletes ^52^. Surprisingly, EGT supplementation alone in cell culture media is sufficient to boost mitochondrial respiration, suggesting that EGT contributes to improved mitochondrial function with exercise training. Proteome-wide thermal shift assays identify MPST as a potential protein target of EGT. *In vitro* binding assays using recombinant MPST confirm that EGT indeed binds MPST. MPST is a sulfurtransferase and one of three mammalian hydrogen sulfide (H_2_S)-producing enzymes, which reacts with various persulfide acceptors, such as thioredoxin, glutathione, L-cysteine or dihydrolipoic acid *in vitro*, to produce H_2_S ^42,43^. However, which molecule is the preferred persulfide acceptor *in vivo* remains unclear and could be context-specific. Two splice variants of MPST give rise to the cytosolic and a dually localized (cytosol and mitochondria) isoform in humans, while rodents only express the shorter, mainly mitochondria localized isoform ^45,46,53^. Consistent with MPST being a direct target of EGT, MPST uses 3-mercaptopyruvate (3-MP) as a substrate to produce pyruvate and H_2_S inside mitochondria, and thereby can contribute to electron flow and mitochondrial respiration; a process important in mouse models of heart failure and obesity ^44,54–59^. While an important role of pyruvate for mitochondrial respiration has long been known, more recently it has been noted, that also H_2_S exhibits multiple positive effects on mitochondrial function. At low concentrations H_2_S is able to boost mitochondrial respiration, while being toxic at high concentrations ^60,61^. 3-MP-derived H_2_S for instance enhances mitochondrial electron transport and bioenergetics through MPST and sulfide quinone oxidoreductase (SQR) at low concentrations, while it is inhibitory at high concentrations ^44^.

Exercise not only increases EGT levels in mitochondria but also increases both mitochondrial localization of MPST and mitochondrial H_2_S production ^62,63^. Our pharmacological and genetic interventions reveal that EGT boosts mitochondrial respiration in an MPST-dependent manner and further support the idea that mitochondrial MPST is a direct target of EGT. Our data are consistent with EGT acting as a small molecule acceptor and reacting with the MPST-bound persulfide to increase pyruvate production and release H_2_S. However, it cannot be excluded that similar to thioredoxin, EGT also acts as an allosteric regulator of MPST or that EGT desulfuration is contributing to H_2_S generation ^64,65^. Alternatively, it is plausible that the ability of MPST to control the activity of a broad range of target proteins involves regulators of mitochondrial respiration and that EGT binding to MPST influences this process ^43^.

While EGT declines in human plasma with age, EGT accumulates in human plasma with exercise and supplementation improves endurance exercise performance in mice ^18,25,29^. Strikingly, increasing the availability of EGT in the diet improves endurance exercise capacity via MPST, a mechanism possibly important for the benefits associated with exercise training. Exercise capacity is a powerful predictor of mortality. An increase in only one metabolic equivalent (1-MET) exercise capacity can confer a 12% improvement in survival ^3^. Given the broad implication of EGT in health and disease, our study provides the basis for future research investigating the role of the EGT-MPST axis in a tissue- and disease-specific manner and during the aging process.

## Acknowledgments

The authors thank all members of the Spiegelman and Sabatini laboratories for valuable discussions and input. The authors would also like to thank the Metabolite Profiling Core Facilities of the Whitehead Institute and Dana-Farber Cancer Institute and Dr. David J. Lefer for sharing the *Mpst*^-/-^ mouse strain with permission of Dr. Noriyuki Nagahara. Schematics were created with BioRender.com. BMS was supported by JPB Foundation 6293803, NIH grants DK123228 and DK119117. HGS was supported by Hope Funds for Cancer Research HFCR-20-03-01-02. MJM was supported by the Deutsche Forschungsgemeinschaft (DFG), Projektnummer 461079553. AA was supported by a Marie-Curie H2020 MSCA Global Fellowship (Grant agreement 101033310). NB was supported by the Deutsche Forschungsgemeinschaft (DFG), Projektnummer 501493132. SDP was supported by Linde Family Foundation, NIBR, 3DC, and DDCF.

## Author contributions

HGS and BMS designed the project and experiments and wrote the manuscript with input from MJM and DMS. MJM performed time course exercise experiments and together with HGS MCK-PGC-1⍺ and mouse phenotyping experiments. YS performed ITC experiments and contributed to OCR measurements. JGVV prepared samples for protein MS and performed data analysis with help from HGS. SS performed NMR experiments and contributed to molecular docking. AJ performed molecular docking. SK performed GTEx analysis. AVC performed replicates of OCR measurements in different cell lines. AMP, JBS and AA contributed to MITO-IP experiments. TZ and BR prepared samples for metabolite MS and performed data analysis with help from HGS. HSS, KS, LS, COY, CB carried out recombinant MPST expression and purification. SDP oversaw MPST expression and purification. AMP and ELM contributed to metabolite MS measurements and analysis. NB, SPG, HA and ETC contributed to data analysis and interpretation.

## Data and materials availability

All data and materials are available from the corresponding authors upon reasonable request. The human plasma metabolomics are from Morville et al., 2020 and can be found at https://www.cell.com/cell-reports/fulltext/S2211-1247(20)31543-6. The human skeletal muscle single cell RNA sequencing data can be found at https://gtexportal.org/home/. The Slc22a4 expression data from primary myotubes are from Ruas et al., 2012 and can be found at https://www.cell.com/cell/fulltext/S0092-8674(12)01363-3. The mass spectrometry data will be made public upon publication.

## Materials and Methods

### Animals

Mice were housed at 22 - 23 °C under a 12 hr light/dark cycle with free access to food and water. MITO-Tag mice (The Jackson Laboratory #032290) were described earlier ^16^ and to generate whole body-expressing MITO-Tag mice, mice were first crossed to mice carrying Cre recombinase under control of the CMV promoter. Afterwards, Cre recombinase was crossed out and whole body-expressing MITO-Tag mice were selected. 8-9 week old male mice were used for the exercise experiments and to isolate mitochondria. Control mice carrying no MITO-Tag (-/-) served as background controls. Hemizygous transgenic MCK-PGC-1⍺ mice were described earlier ^36^. 8 week old male transgenic mice were used for tissue harvesting and wild-type littermates served as controls. For treadmill experiments 8 week old male mice were used (C57BL/6, The Jackson Laboratory #000664). For voluntary wheel running and ergothioneine-diet experiments 8-11 week old male control (C57BL/6, The Jackson Laboratory #000664) and *Mpst^-/-^* ^54,66^ (C57BL/6) mice were used. For ergothioneine quantification in gastrocnemius muscle tissue 15-16 week old male (C57BL/6, The Jackson Laboratory #000664) mice were used. Animal experiments were performed according to procedures approved by the Institutional Animal Care and Use Committee (IACUC) of the Massachusetts Institute of Technology and the Beth Israel Deaconess Medical Center.

### Exercise training protocols and diet feeding experiments

For MITO-IP experiments mice were single housed and half of them were placed in cages containing an in-cage running wheel for 4 weeks (Starr Life Sciences). Distance run was continuously recorded. Body weight and grip strength were assessed in both, trained and untrained mice. Running wheels were removed 4 - 6 hours before tissue harvesting to avoid metabolic differences due to acute exercise performance.

To assess exercise performance upon ergothioneine-enriched diet feeding, mice were single housed and received a control diet (Research Diets Product# D11112201) or an ergothioneine-enriched diet (Research Diets Product# D22021803, Ergothioneine from Thermo Scientific AAJ6786103) for 2 weeks. After 2 weeks feeding, mice were placed in cages containing an in-cage running wheel for 6 weeks (Starr Life Sciences). Distance run was continuously recorded. Body weight and grip strength were assessed in both groups at the end of the experiment. Throughout the recording mice kept *ad libitum* access to the respective diet. To determine body weight and food consumption during the first 2 weeks of feeding, mice were single housed and received the control diet or the ergothioneine-enriched diet while continuously recording body weight and food consumption. For the acute exercise time course experiments mice were trained for 5 min at 12 m/min followed by 1 min rest. Subsequently, mice run 5 min at 12 m/min and 5 min at 14 m/min. On the third day of training, sedentary mice were removed from the treadmill after training and exercise mice were kept running for a total of 45 min with ramped up speed of 2 m/min every 5 min and maximum speed of 26 m/min. Tissues were collected 0 min, 30 min, 60 min, 2 h or 4 h after the run ^12^.

### Rapid isolation of mitochondria and metabolite extraction from gastrocnemius muscle tissue and cultured cells

Isolation of mitochondria was performed as previously described with modifications ^16^. All procedures were performed on ice or in a 4 °C cold room. The entire gastrocnemius from MITO-Tag Mice was excised and minced quickly on an ice-cold metal block using a razor blade. Afterwards, the tissue was homogenized in 1 ml of KPBS (136 mM KCl, 10 mM KH_2_PO_4_, pH 7.25, in LC/MS water) with 25 strokes using a tissue grinder with glass pestle. The homogenate was spun down at 1000 x g for 2 min at 4 °C. 25 µl of the supernatant was taken as a sample of gastrocnemius tissue and was extracted in the appropriate reagent depending on the downstream analysis. The same procedure was applied to isolate metabolites from gastrocnemius muscle in MCK-PGC-1⍺ and mouse diet experiments using ice-cold 80:20 methanol:water with internal standards. In the case of MITO-Tag Mice the remaining supernatant was subjected to immunoprecipitation with prewashed magnetic anti-HA beads for 3.5 min, followed by three washes in 1 ml KPBS. In the final wash 150 µl of the suspension of beads was set aside and used for protein extraction and immunoblotting by aspirating the KPBS and resuspending the beads in 50 µl of ice-cold Triton lysis buffer. The remainder of the beads were then resuspended in 50 µl ice-cold 80:20 methanol:water with internal standards.

To isolate metabolites from cultured cells, medium was removed, cells were washed with ice-cold PBS, and metabolites were extracted in ice-cold 80:20 methanol:water with internal standards (Sigma Aldrich, 909653) for 15 min at -20 °C. Plates were scraped on dry ice and lysates collected, vortexed and centrifuged twice at 17000 x g for 5 min at 4 °C. Supernatants were collected and stored at -80 °C until further processing.

### Metabolomics MITO-IP and MCK-PGC-1⍺ experiments

Metabolite profiling was conducted on a QExactive bench top orbitrap mass spectrometer equipped with an Ion Max source and a HESI II probe, which was coupled to a Dionex UltiMate 3000 HPLC system (Thermo Fisher Scientific, San Jose, CA). External mass calibration was performed using the standard calibration mixture every 7 days. Typically, 2 µl sample were injected onto a SeQuant® ZIC®-pHILIC 150 x 2.1 mm analytical column equipped with a 2.1 x 20 mm guard column (both 5 mm particle size; EMD Millipore). Buffer A was 20 mM ammonium carbonate, 0.1% ammonium hydroxide; Buffer B was acetonitrile. The column oven and autosampler tray were held at 2 °C and 4 °C, respectively. The chromatographic gradient was run at a flow rate of 0.150 mL/min as follows: 0-20 min: linear gradient from 80-20% B; 20-20.5 min: linear gradient form 20-80% B; 20.5-28 min: hold at 80% B. The mass spectrometer was operated in full-scan, polarity-switching mode, with the spray voltage set to 3.0 kV, the heated capillary held at 275 °C, and the HESI probe held at 350 °C. The sheath gas flow was set to 40 units, the auxiliary gas flow was set to 15 units, and the sweep gas flow was set to 1 unit. MS data acquisition was performed in a range of *m/z* = 70–1000, with the resolution set at 70,000, the AGC target at 1x10^6^, and the maximum injection time at 20 msec. Relative quantitation of polar metabolites was performed with TraceFinder™ 4.1 (Thermo Fisher Scientific) using a 5 ppm mass tolerance and referencing an in-house library of chemical standards. All solvents, including water, were purchased from Fisher and were Optima LC/MS grade.

### Metabolomics cell culture, mouse diet experiments and ergothioneine quantification

Targeted ergothioneine analysis was conducted on a QExactive HF-X mass spectrometer. The analytes were ionized via a HESI II probe. The mass spectrometer was coupled to a Vanquish binary UPLC system (Thermo Fisher Scientific, San Jose, CA). For chromatographic separation prior to mass analysis, 5 μL of the sample was injected onto a BEH Amide (50 mm length) or BEH Z-HILIC column (100 mm length) (both with 1.7 µm particle size, 2.1 mm internal diameter, Waters). The method was adapted from Mülleder et al ^67^. Both mobile phases were buffered with 10 mM ammonium formate and 45 mM formic acid (pH 2.7). In mobile phase A, the buffers were in 1:1 acetonitrile:water. Mobile phase B consisted of buffer in 95:5:5 acetonitrile:water:methanol. The column oven was held at 45 °C and autosampler at 4 °C. The chromatographic gradient was run at a flow rate of 0.4 ml/min as follows: 0.75 min initial hold at 95% B; 0.75-3.00 min linear gradient from 95% to 30% B, 1.00 min isocratic hold at 30% B. B was brought back to 95% over 0.50 minutes, after which the column was re-equilibrated under initial conditions. The mass spectrometer was operated in full-scan or PRM positive mode, with the spray voltage set to 3.5 kV, the capillary temperature to 320 °C, and the HESI probe to 300 °C. The sheath gas flow was set to 50 units, the auxiliary gas flow was set to 10 units, and the sweep gas flow was set to 1 unit. MS^1^ data acquisition was performed in a range of *m/z* = 75–1050, with the resolution set at 120,0000. MS^2^ data were acquired at resolution 30,000. Blank samples were injected at regular intervals. Retention times and fragmentation patterns were determined using authentic standards. For full scan quantitation, M^+^ ion at *m/z* 230.0963 was used, and for PRM data, a characteristic fragment was detected at *m/z* 186.1049; a 3 mmu tolerance was allowed in the measured *m/z*. Raw data were converted to .mzML and processed using eMZed ^68^ as follows. For relative quantification, baseline was determined and subtracted from ergothioneine peak area. The corresponding blank signal was then subtracted from the ergothioneine peak area. Biomass normalization was performed by dividing the ergothioneine peak area by the peak area of endogenous glutamate. Lastly, the ergothioneine peak area was normalized to technical variability by dividing ergothioneine peak area by the nearest coeluting amino acid in the internal standard (typically arginine). For absolute quantification, a triplicate of log2 dilutions of ergothioneine were prepared from 16 nM to 16 µM and a standard curve was fit by linear regression using LinearRegression function from sklearn. Each sample was confirmed to lie within the linear range of the standard curve. Water, acetonitrile, methanol, and formic acid were purchased from Fisher and were Optima LC/MS grade. Ammonium formate powder was purchased from Honeywell Fluka.

### Western blotting

Samples were lysed in Triton lysis (50 mM Tris-HCl, pH7.4, 150 mM NaCl, 1 mM EDTA, 1% (vol/vol) Triton X-100) or RIPA buffer (Cell Signaling) and cleared by centrifugation. Protein quantification was performed using Pierce BCA Protein Assay Kit (Thermo Scientific) and proteins were separated on NuPAGE 4-12% Bis-Tris Gels (Invitrogen). Proteins were subsequently transferred to a 0.45 µm PVDF membrane (Millipore). Primary antibodies were used at the following dilutions in 5% Blotting-Grade Blocker (BioRad): VDAC (1:1000, Cell Signaling, rabbit, #4661) RRID: AB_10557420, SDHA (1:1000, Proteintech, rabbit, #14865-1-AP) RRID: AB_11182164, Citrate Synthase (CS, 1:1000, Cell Signaling, rabbit, #14309) RRID:AB_2665545, Cathepsin B (CTSB, 1:1000, Cell Signaling, rabbit, #31718) RRID: AB_2687580, Golgin 97 (1:1000, Cell Signaling, rabbit, #13192) RRID: AB_2798144, GAPDH (1:1000, Cell Signaling, rabbit, #5174) RRID: AB_10622025, Calreticulin (CALR, 1:1000, Cell Signaling, rabbit, #12238) RRID: AB_2688013, PEX13 (1:1000, Millipore, rabbit, #ABC143) https://www.emdmillipore.com/US/en/product/Anti-PEX13-peroxisomal-membrane-marker-Antibody, MM_NF-ABC143, SLC22A4 (1:500, Abnova, mouse, #H00006583-A01) RRID: AB_627475, OXPHOS cocktail (1:1000, Abcam, mouse, #ab110413) RRID: AB_2629281, MPST (1:1000, Abcam, rabbit, #ab154514) RRID: AB_2895038. Secondary antibodies (Anti-Rabbit/Mouse IgG HRP Conjugate, Promega) were used at 1:5000 in 5% Blotting-Grade Blocker (BioRad).

### Gene expression analysis by quantitative reverse transcription PCR

Total RNA was isolated using TRIzol (Invitrogen 15596018) and RNeasy Mini purification kit (Qiagen 74104) according to manufactures protocols. DNA was digested on column using RNase-Free DNase Set (Qiagen 79254). RNA was reverse transcribed using High-Capacity cDNA Reverse Transcription kit with RNase Inhibitor (Applied Biosystems 4374966) and gene expression was determined by quantitative PCR (QuantStudio 6 Pro Real-Time PCR System) with GoTag qPCR System (Promega, A6002). *Actb* and *Rplp0* were used as housekeeping genes. The following primer sequences were used: mouse *Actb* (IDT assay ID: Mm.PT.39a22214843.g), mouse *Rplp0* (fwd: AGATTCGGGATATGCTGTTGGC, rev: TCGGGTCCTAGACCAGTGTT), mouse *Ppargc1a* (1) (IDT assay ID: Mm.PT.58.17390716), mouse *Ppargc1a* (2) (fwd: GGTTGGTGAGGACCAGCC, rev: AATCCACCCAGAAAGCTGTCT), mouse *Slc22a4* (IDT assay ID: Mm.PT.58.31033975), mouse *Myh7* (fwd: CTACAGGCCTGGGCTTACCT, rev: TCTCCTTCTCAGACTTCCGC), mouse *Myh2* (fwd: ATCCAAGTTCCGCAAGATCC, rev: TTCGGTCATTCCACAGCATC), mouse *Myh1* (fwd: ATGAACAGAAGCGCAACGTG, rev:AGGCCTTGACCTTTGATTGC), mouse *Myh4* (fwd: AGACAGAGAGGAGCAGGAGAGTG , rev: CTGGTGTTCTGGGTGTGGAG), mouse *Mpst* (IDT assay ID: Mm.PT.58.5446715).

### Human Genotype Expression (GTEx) Project data analysis

The Genotype-Tissue Expression (GTEx) Project was supported by the Common Fund of the Office of the Director of the National Institutes of Health, and by NCI, NHGRI, NHLBI, NIDA, NIMH, and NINDS. The data used for the analyses described in this manuscript were obtained from the GTEx Portal on 03/03/2022.

### Cell culture

To obtain primary inguinal white adipocytes the inguinal white adipose stromal-vascular fraction (SVF) was isolated from 6-10 week old wild-type mice. In detail, iWAT was dissected, washed with ice-cold HBSS, and minced. Subsequently, suspension was incubated in HBSS (Life Technologies, 14025-092) containing 10 mg/ml collagenase D (Sigma Aldrich, 11088882001), 3 U/ml dispase II (Roche Diagnostics, 4942078001), and 10 mM CaCl_2_ for 30 min at 37 °C with occasional shaking. To stop collagenase reaction, complete adipocyte culturing medium (DMEM/F-12 GlutaMAX (Life Technologies, 10565042), 10% fetal bovine serum (BenchMark,100-106), 1X PenStrep (Life Technologies, 15140122), 100 mg/ml Primocin (Fisher Scientific, NC9141851)) was added to the suspension and filtered through a 100 mm cell strainer. Next, cell suspension was centrifuged at 600 x g for 5 min, and SVF pellet was resuspended in complete adipocyte culturing medium, filtered through a 40 mm cell strainer and centrifuged at 600 x g for 5 min. SVF pellet was resuspended and plated in complete adipocyte culturing medium. Cells were split two times at a 1:3 ratio when confluency reached 70%. For experiments, cells were grown until confluency and differentiation of pre-adipocytes was induced by treatment with an adipogenic cocktail (1 µM rosiglitazone (Cayman Chemical 71740), 0.5 mM 3-Isobutyl-1-methylxanthine (IBMX) (Sigma I5879), 1 µM dexamethasone (Sigma D4902), 870 nM insulin (Sigma I5500)) in complete adipocyte culture medium for 2 days. Subsequently, medium was changed to complete adipocyte maintenance medium with 1 µM rosiglitazone and 870 nM insulin for 6 days.

C2C12 myoblasts and HeLa cells were cultured in DMEM (Corning 10-017-CV) supplemented with 10% fetal bovine serum and 1% penicillin/streptomycin. To differentiate C2C12 myoblasts into myotubes, myoblasts were cultured in DMEM (Corning 10-017-CV) supplemented with 2 % donor horse serum and 1% penicillin/streptomycin for 3 days. To generate *MPST^-/-^* HeLa cells single guide RNAs (sgRNAs) against *MPST* (fwd: CACCGGCGTCGTAGATCACGACGT, rev: AAACACGTCGTGATCTACGACGCC) were subcloned into the plentiCRISPR V1 (Addgene 52963). Subcloned plasmids were co-transfected into HEK293T cells with lentiviral packaging vectors. HeLa cells were subsequently infected with the lentivirus and selected with puromycin, generating stable knockout cell lines. Limiting dilution was used to isolate single cell clones. Clones were cultured and screened for the relevant knockouts by western blotting. The parental cell line and a single clone still expressing MPST after limiting dilution were used as control cell lines. All cells were incubated at 37 °C and 5% CO_2_.

### Mitochondrial respiration

Mitochondrial respiration was determined using the XF24 Extracellular Flux Analyzer (Seahorse Bioscience). Differentiated primary inguinal adipocytes (primary iWAT) were counted and equal amounts seeded on seahorse plates. Cells were treated with vehicle or 500 µM ergothioneine (Sigma E7521) and incubated at 37 °C and 5% CO_2_ for 72 h. 500 nM norepinephrine (Sigma A9512) was used to induce norepinephrine-stimulated respiration. Uncoupled and maximal respiration was determined using 5 µM oligomycin (Sigma 495455) and 5 µM FCCP (Sigma C2920). 5 µM rotenone (Sigma R8875) and 5 µM antimycin A (Sigma A8674) were used to inhibit complex I- and complex III-dependent respiration. C2C12 myoblasts were differentiated into myotubes in seahorse plates as described above in the presence of vehicle 10 or 500 µM ergothioneine during the 3-day differentiation period. To inhibit MPST, cells were pretreated with 100 - 200 µM I3MT-3 (Selleck S3541) for 3 - 4 h before measuring respiration. Oligomycin, FCCP, rotenone and antimycin A were all used at a final concentration of 2 µM. Equal amounts of control and *MPST^-/-^* HeLa cells seeded on seahorse plates. Cells were treated with vehicle or 500 µM ergothioneine and incubated at 37 °C and 5% CO_2_ for 72 h. Oligomycin, FCCP, rotenone and antimycin A were all used at a final concentration of 2 µM.

### Lysate-based proteome integral solubility alteration (PISA)

Differentiated primary inguinal adipocytes (primary iWAT) were suspended in lysis buffer (1X PBS, 1.5 mM MgCl_2_, 1X protease inhibitor (Pierce protease inhibitor mini tablets) and the proteomes were extracted using a dounce homogenizer. The resulting lysate was spun at 300 x g for 3 min to remove any unbroken cells. The lysate was diluted to 2 mg/mL using lysis buffer and allowed to warm to room temperature. Ergothioneine (solubilized in water) or water was added to the desired concentration (50 µM, 500 µM, or 1 mM) and the samples were allowed to incubate at room temperature for 30 min with gentle shaking. Each sample was divided into ten aliquots, each of which was heated to a different temperature from 48 - 58 °C for 3 min in an Eppendorf Mastercycler Pro S. Samples were allowed to cool at room temperature for 5 min. An equal volume of ice cold 2X extraction buffer (1X PBS, 1.5 mM MgCl_2_, 1% NP-40, 1X protease inhibitor) was added to each PCR tube to achieve a final NP-40 concentration of 0.5%. The samples were allowed to incubate at 4 °C for 10 min. An equal volume of each sample was pooled and spun at 21,000 x g for 90 min. ∼15 μg of soluble protein were collected and prepared for LC-MS/MS analysis.

### Proteomics LC-MS/MS sample preparation

Samples (15-20 μg protein primary iWAT) were diluted in prep buffer (400 mM EPPS pH 8.5, 1% SDS, 10 mM TCEP) and incubated at room temperature for 10 min. Iodoacetimide was added to a final concentration of 10 mM to each sample and incubated for 25 min in the dark. Finally, DTT was added to each sample to a final concentration of 10 mM. A buffer exchange was carried out using a modified SP3 protocol ^69,70^. Briefly, ∼250 μg of each SpeedBead Magnetic Carboxylate modified particles (Cytiva; 45152105050250, 65152105050250) mixed at a 1:1 ratio were added to each sample. 100% ethanol was added to each sample to achieve a final ethanol concentration of at least 50%. Samples were incubated with gentle shaking for 15 min. Samples were washed three times with 80 % ethanol. Protein was eluted from SP3 beads using 200 mM EPPS pH 8.5 containing trypsin (ThermoFisher Scientific) and Lys-C (Wako). Samples were digested overnight at 37 °C with vigorous shaking. Acetonitrile was added to each sample to achieve a final concentration of 30%. Each sample was labelled, in the presence of SP3 beads, with ∼65 μg of TMTpro-16plex reagents (ThermoFisher Scientific) ^71,72^. Experimental layouts for each experiment were described in corresponding source data tables. Following confirmation of satisfactory labelling (>97%), excess TMTpro reagents were quenched by addition of hydroxylamine to a final concentration of 0.3%. The full volume from each sample was pooled and acetonitrile was removed by vacuum centrifugation for one hour. The pooled sample was acidified using formic acid and peptides were de-salted using a Sep-Pak Vac 50 mg tC18 cartridge (Waters). Peptides were eluted in 70% acetonitrile, 1% formic acid and dried by vacuum centrifugation. The peptides were resuspended in 10 mM ammonium bicarbonate pH 8, 5% acetonitrile and fractionated by basic pH reverse phase HPLC. In total 24 fractions were collected. The fractions were dried in a vacuum centrifuge, resuspended in 5% acetonitrile, 1% formic acid and desalted by stage-tip. Final peptides were eluted in, 70% acetonitrile, 1% formic acid, dried, and finally resuspended in 5% acetonitrile, 5% formic acid. In the end, 12 of 24 fractions were analyzed by LC-MS/MS.

### Proteomics mass spectrometry data acquisition

Data were collected on an Orbitrap Fusion Lumos mass spectrometer (ThermoFisher Scientific) coupled to a Proxeon EASY-nLC 1000 LC pump (ThermoFisher Scientific). Peptides were separated using a 180-min gradient at 500 nL/min on a 30-cm column (i.d. 100 μm, Accucore, 2.6 μm, 150 Å) packed inhouse. MS1 precursor scans were acquired in the orbitrap at 120 K resolution, 1e6 AGC target with a maximum of 50 ms injection time. Data dependent, “top 10” MS2 scans were acquired in the ion trap with collisional induced dissociation (CID) fragmentation. While setting varied from instrument to instrument and with instrument performance, generally the following was used for MS2 scans: NCE 35%, 1.2e4 AGC target, maximum injection time 50 ms, isolation window 0.5 Da. Orbiter, an on-line real-time search algorithm, was used to trigger MS3 quantification scans ^73^. MS3 scans were acquired in the orbitrap using varying settings based on instrument performance: 50,000 resolution, AGC of 2.5 × 10^5^, injection time of 200 ms, HCD collision energy of 65%. Protein-level closeout was typically set to two peptides per protein per fraction.

### Proteomics mass spectrometry data analysis

Raw files were first converted to mzXML, and monoisotopic peaks were re-assigned using Monocle ^74^. Database searching included all mouse entries from Uniprot (downloaded in July, 2014). The database was concatenated with one composed of all protein sequences in the reversed order. Sequences of common contaminant proteins (e.g., trypsin, keratins, etc.) were appended as well. Searches were performed using the comet search algorithm. Searches were performed using a 50-ppm precursor ion tolerance and 1.0005 Da fragment ion tolerance. TMTpro on lysine residues and peptide N termini (+304.2071 Da) and carbamidomethylation of cysteine residues (+57.0215 Da) were set as static modifications, while oxidation of methionine residues (+15.9949 Da) was set as a variable modification. Peptide-spectrum matches (PSMs) were adjusted to a 1% false discovery rate (FDR) ^75^. PSM filtering was performed using linear discriminant analysis (LDA) as described previously ^76^, while considering the following parameters: comet log expect, different sequence delta comet log expect (percent difference between the first hit and the next hit with a different peptide sequence), missed cleavages, peptide length, charge state, precursor mass accuracy, and fraction of ions matched. Each run was filtered separately. Protein-level FDR was subsequently estimated at a data set level. For each protein across all samples, the posterior probabilities reported by the LDA model for each peptide were multiplied to give a protein-level probability estimate. Using the Picked FDR method ^77^, proteins were filtered to the target 1% FDR level. For reporter ion quantification, a 0.003 Da window around the theoretical *m/z* of each reporter ion was scanned, and the most intense *m/z* was used. Reporter ion intensities were adjusted to correct for the isotopic impurities of the different TMTpro reagents according to manufacturer specifications. Peptides were filtered to include only those with a summed signal-to-noise (SN) of 160 or greater across all channels. For each protein, the filtered peptide TMTpro SN values were summed to generate protein quantification.

Statistical analysis was performed using Perseus ^78^. *P*-values were calculated using Student’s t-test. Fold changes were calculated by averaging abundance of each group and dividing “treated” group by control group average. Q-values were calculated by a permutation-based false discovery rate estimation and proteins with q-values < 0.05 were considered statistically significant.

### Protein expression and purification recombinant MPST

The human MPST expression plasmid with the N-terminal His tag with TEV cleavage site (MPSTA) was a gift from Nicola Burgess-Brown (Addgene plasmid #42482). The N-terminal His tag construct of human MPST was co-transformed with GroES-EL chaperon plasmid from Takara (#3340), overexpressed in *E. coli* BL21 (DE3) and purified using affinity chromatography and size-exclusion chromatography. Briefly, cells were grown at 37 °C in TB medium in the presence of 50 μg/mL of kanamycin, 20 μg/mL chloramphenicol, and 0.5 mg/mL L-arabinose to an OD of 0.8, cooled to 17 °C, induced with 100 μM isopropyl-1-thio-D-galactopyranoside (IPTG), incubated overnight at 17 °C, collected by centrifugation, and stored at -80 °C. Cell pellets were lysed in buffer A (25 mM HEPES, pH 7.5, 500 mM NaCl, and 0.5 mM TCEP) using Microfluidizer (Microfluidics), and the resulting lysate was centrifuged at 30,000 x g for 40 min. Ni-NTA beads (Qiagen) were mixed with cleared lysate for 90 min and washed with buffer A. Beads were transferred to an FPLC-compatible column, and the bound protein was washed further with buffer A for 10 column volumes and eluted with buffer B (25 mM HEPES, pH 7.5, 500 mM NaCl, 10% glycerol, 0.5 mM TCEP, and 400 mM imidazole). The eluted sample was concentrated and purified further using a Superdex 200 16/600 column (Cytiva) in buffer C containing 20 mM HEPES, pH 7.5, 300 mM NaCl, and 0.5 mM TCEP. SEC fractions, containing MPST, were concentrated to ∼24 mg/mL and stored at -80 °C.

### Nuclear magnetic resonance spectroscopy (NMR)

The human MPST expression plasmid with the N-terminal His tag with TEV cleavage site (MPSTA) was a gift from Nicola Burgess-Brown (Addgene plasmid #42482). This construct was co-expressed with chaperones groEL, groES, and tig (pG-Tf2 plasmid, Takara #3340). 1 L of M9 minimal medium supplemented with 1 g ^15^NH_4_Cl was inoculated to an OD600 of 0.05 with a LB overnight culture and grown to an OD600 of 0.8 in the presence of chloramphenicol (20 μg/L) and kanamycin (50 μg/L). The culture was cooled to 17 °C for 45 min and expression of chaperones and MPST was induced with 10 μg/L tetracycline and 0.1 mM IPTG, respectively. Cells were incubated for 24 h at 17 °C, harvested by centrifugation, and resuspended in lysis buffer (25 mM HEPES pH 7.5, 500 mM NaCl, 20 mM imidazole, 0.5 mM TCEP, 10% glycerol) supplemented with lysozyme, protease inhibitor, and benzonase. Cells were then lysed by sonication and the lysate was centrifuged at 30,000 x g for 45 min. His-tagged protein was bound to Ni-NTA beads (Qiagen). The beads were washed with 100 mL lysis buffer and protein was eluted with 40 mL elution buffer (25 mM HEPES pH 7.5, 500 mM NaCl, 300 mM imidazole, 0.5 mM TCEP, 10% glycerol). The N-terminal His tag was cleaved off by incubation with TEV protease overnight at 4 °C in a buffer containing 25 mM HEPES pH 7.5, 300 mM NaCl, 0.5 mM TCEP, and 0.5 mM EDTA. Protein was concentrated and further purified using a Superdex 200 16/600 column (Cytiva) into NMR buffer (50 mM sodium phosphate pH 7.0, 100 mM NaCl, 0.5 mM TCEP).

^1^H^15^N HSQC spectra were collected using 50 μM or 600 μM protein in NMR buffer supplemented with 5% D_2_O. Ergothioneine, 3-mercaptopyruvate, glutathione and L-cysteine were dissolved in NMR buffer to generate stock solutions at 20 mM. When required, appropriate amount of the stock solutions was added to the protein-containing samples to reach a final concentration of 500 μM (for the 50 μM protein samples) or 1.2 mM (for the 600 μM protein sample). ^1^H^15^N-HSQC spectra of the 600 μM samples were recorded on a Bruker Avance III 800 MHz spectrometer equipped with a TXO-style cryogenically cooled probe. 64 scans and 128 complex points in the indirect ^15^N dimension were collected at 298 K. The indirect dimension was sampled using an Echo-AntiEcho acquisition mode. ^1^H^15^N-HSQC spectra of the 50 μM samples were recorded on a Bruker Ascend 600 MHz spectrometer equipped with a QCI-style cryogenically cooled probe. 256 scans and 128 complex points in the indirect ^15^N dimension were collected at 298 K. The indirect dimension was sampled using an Echo-AntiEcho acquisition mode. NMR data was processed using nmrPipe ^79^ using the same script for all the spectra derived from the same magnet. Spectra were then further analyzed using the ccpNMR software (version 3.2.0) ^80^.

### Isothermal titration calorimetry (ITC)

Recombinant human MPST was dialyzed against Tris buffer (20 mM Tris pH 8.0, 300 mM NaCl, 0.5 mM TCEP). Ergothioneine (Sigma E7521) was dissolved in the same buffer. ITC experiments were carried out in an Affinity ITC instrument (TA Instruments) at 25 °C. The titrations were performed by injecting 2.5 µL aliquots of 500 µM ergothioneine into the calorimeter cell containing a 185 µL solution of 27.5 µM MPST with a constant stirring speed at 125 rpm and heat was recorded. The data were analyzed with NanoAnalyze using the independent fit model. All the uncertainties were estimated by the native statistics module with 10000 synthetic trials and 95% confidence level.

### Structural modeling of MPST and ergothioneine

Amino acid sequence of 3-mercaptopyruvate sulfurtransferase was retrieved from UniProt ^81^ (UniProt ID: P25325). Sequence 11 to 288 of the 297 amino acid sequence of 3-mercaptopyruvate sulfurtransferase was found to be modellable using PDB structure 4JGT. The modified cysteine in the active site was kept as S-mercaptocysteine (position: 248). The structure was prepared for docking using Maestro module of Schrodinger software tool, Schrödinger Release 2023-4: Maestro, Schrödinger, LLC, New York, NY, 2023. 3-mercaptopyruvate sulfurtransferase was then docked to ergothioneine using Glide ^82^ along with nine variants of AutoDock (Vina, Smina, Qvina, Unidock, QvinaW, VinaXB, Gwovina, Psovina and QvinaW_serial) ^83–90^ to perform consensus docking.

### Statistical analyses

Data were expressed as box and whisker plots with center lines denoting medians; box limits denote 25^th^ and 75^th^ percentiles; whiskers denote maxima and minima or mean ± s.e.m. *P* values were calculated using two-tailed Student’s *t*-test for pairwise comparison of variables, one-way ANOVA for multiple comparison of variables, and two-way ANOVA for multiple comparison involving two independent variables.

---

**Fig. S1.**
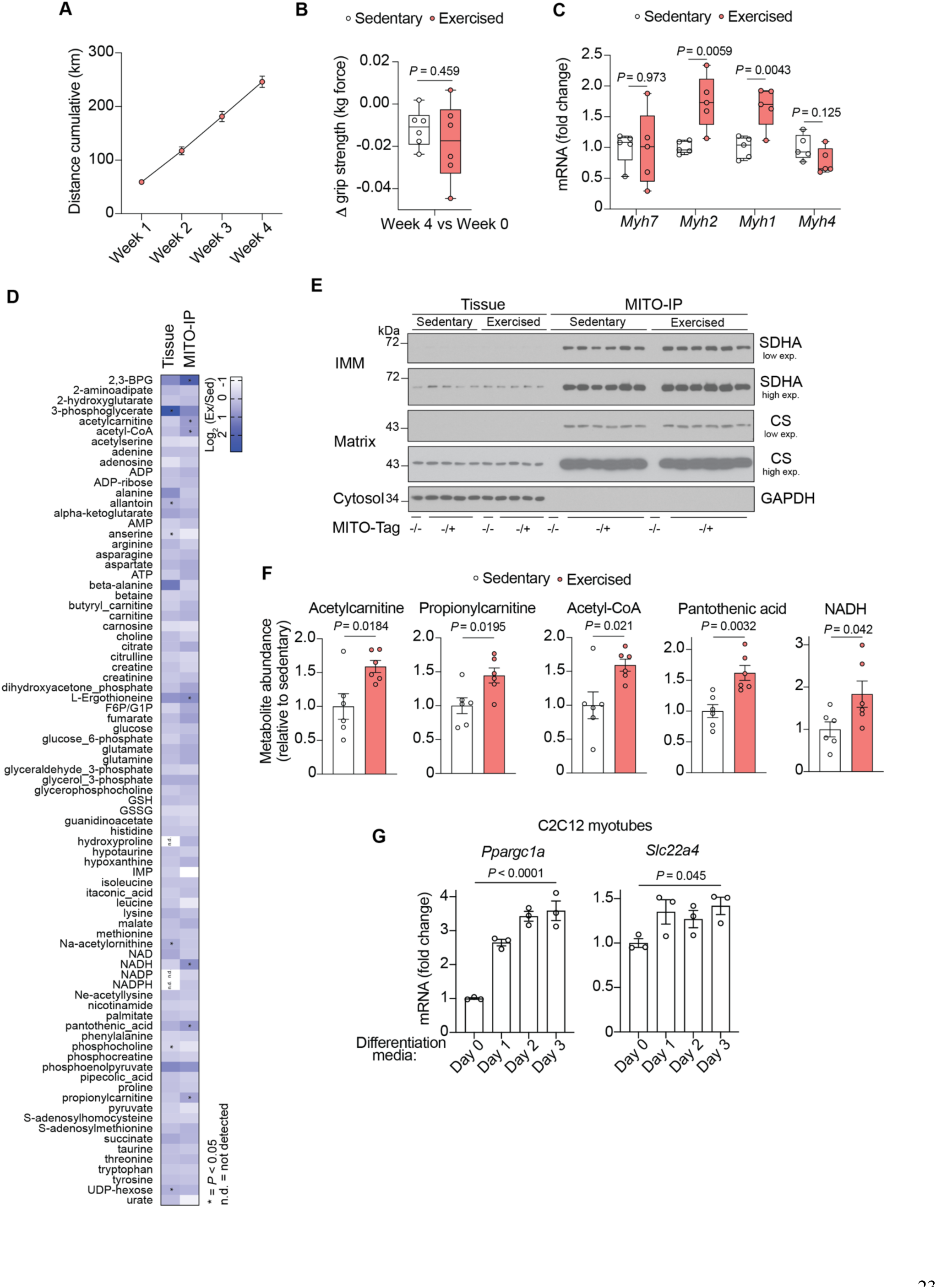
MITO-IP exercised muscle and *Slc22a4* regulation. (**A**) Cumulative running distance MITO-Tag Mice (-/+, n = 6) during 4 weeks of voluntary wheel running (VWR). (**B**) Change in grip strength (kg force) comparing timepoint before VWR (week 0) and after 4 weeks VWR (week 4) (n = 6). (**C**) Fiber type marker expression in gastrocnemius muscle from sedentary and exercised mice after 4 weeks of VWR monitored by RT-qPCR (n = 5). (**D**) Heat map of changes in metabolites as assessed at the mitochondrial and tissue level (gastrocnemius muscle) in sedentary and exercised (4 weeks VWR) MITO-Tag Mice (-/+, n = 6). The data are presented as the log_2_-transformed mean fold difference (Exercised/Sedentary). Metabolites included in this heat map fulfilled criteria to be mitochondrial as described previously ^16^. N.d. = not detected, * = *P* < 0.05. (**E**) Immunoblot analysis of gastrocnemius muscle tissue and the anti-HA immunoprecipitates (MITO-IP) from background control mice (-/-, n = 1) and mice expressing one copy of the MITO-Tag (-/+, n = 6. For tissue samples only four representative mice are shown per condition due to limited number of gel slots available). IMM, inner mitochondrial membrane; Matrix, mitochondrial matrix. low exp. = low exposure, high exp. = high exposure. (**F**) Comparison of metabolites significantly changed in exercised (-/+, n = 6) vs. sedentary (-/+, n = 6) anti-HA immunoprecipitates (MITO-IP) from gastrocnemius muscle. (**G**) *Ppargc1a* and *Slc22a4* expression in C2C12 cells at different timepoints during differentiation from myoblasts into myotubes (n = 3) monitored by RT-qPCR. Data are means ± s.e.m. (A, F, G). Centre lines denote medians; box limits denote 25^th^ and 75^th^ percentiles; whiskers denote maxima and minima (B, C). *P* values calculated using two-tailed unpaired t-test (B, C, D, F), one-way analysis of variance (ANOVA) (G).

**Fig. S2.**
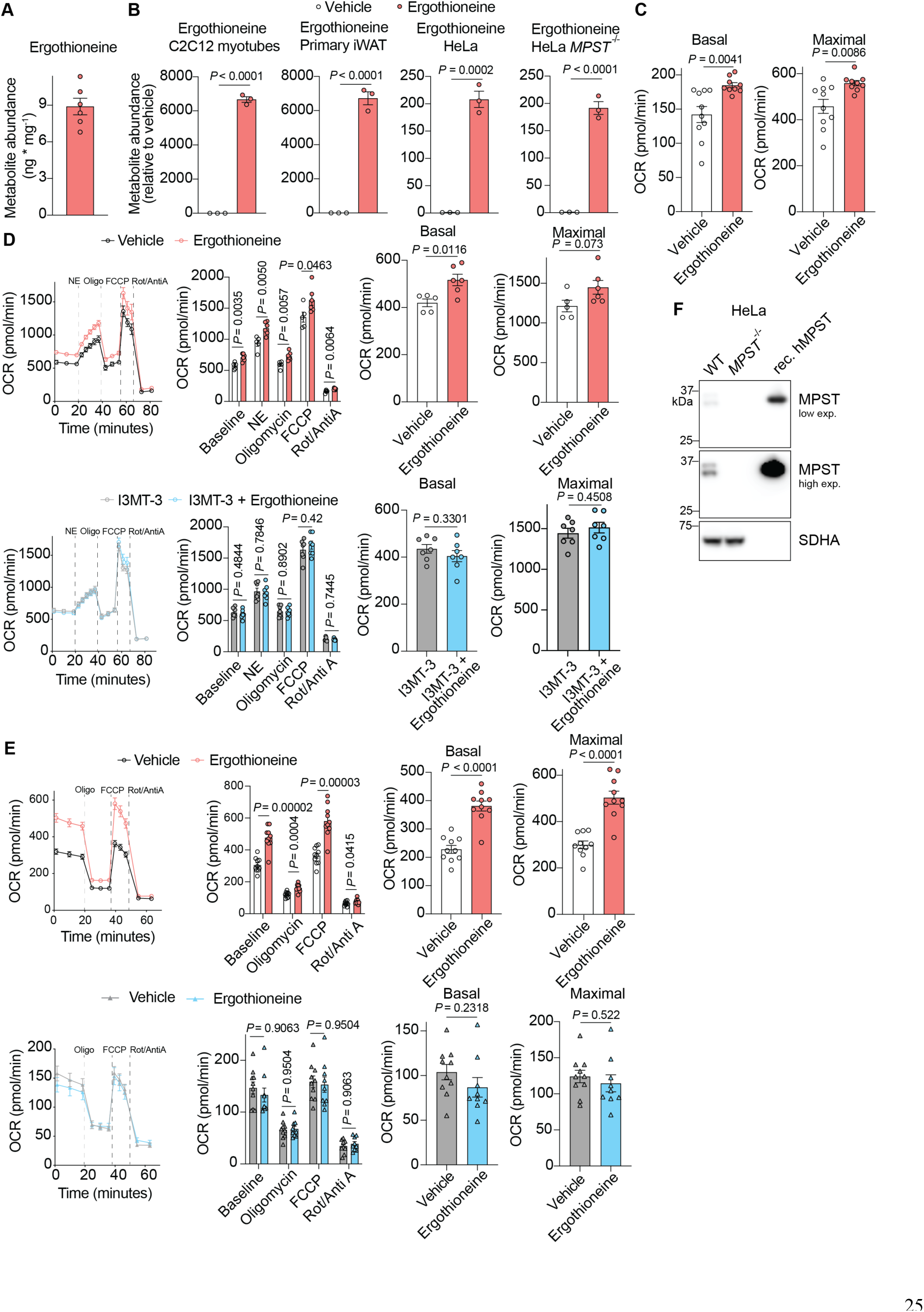
Ergothioneine levels and mitochondrial respiration. (**A**) Ergothioneine levels in gastrocnemius muscle tissue from sedentary mice (n = 6). (**B**) Ergothioneine levels in cells treated with vehicle (n = 3) or 500 µM ergothioneine (n = 3) for 72 h. (**C**) Basal and maximal mitochondrial respiration of C2C12 myotubes. Equal amounts of cells were treated with vehicle (n = 10) or 500 µM ergothioneine (n = 9) for 72 h. (**D**) Oxygen consumption rates (OCR) of primary iWAT cells after treatment with norepinephrine (NE), oligomycin (Oligo), carbonyl cyanide p-triflouromethoxyphenylhydrazone (FCCP), rotenone (Rot), antimycin A (AntiA). Equal amounts of cells were treated with vehicle (n = 5) or 500 µM ergothioneine (n = 6) for 72 h (upper panels). For I3MT-3 treatment cells were pre-treated with I3MT-3 for 3 h before the measurement (I3MT-3, n = 7; I3MT-3 + ergothioneine, n = 7) (lower panels). (**E**) OCRs of HeLa cells after treatment with Oligo, FCCP, Rot, AntiA. Equal amounts of cells were treated with vehicle (WT upper panel, n = 10; *MPST^-/-^* lower panel, n = 10) or 500 µM ergothioneine (WT upper panel, n = 10; *MPST^-/-^* lower panel, n = 9) for 72 h. (**F**) Immunoblot analysis of HeLa cells. Data are means ± s.e.m.. *P* values calculated using two-tailed unpaired t-test.

**Fig. S3.**
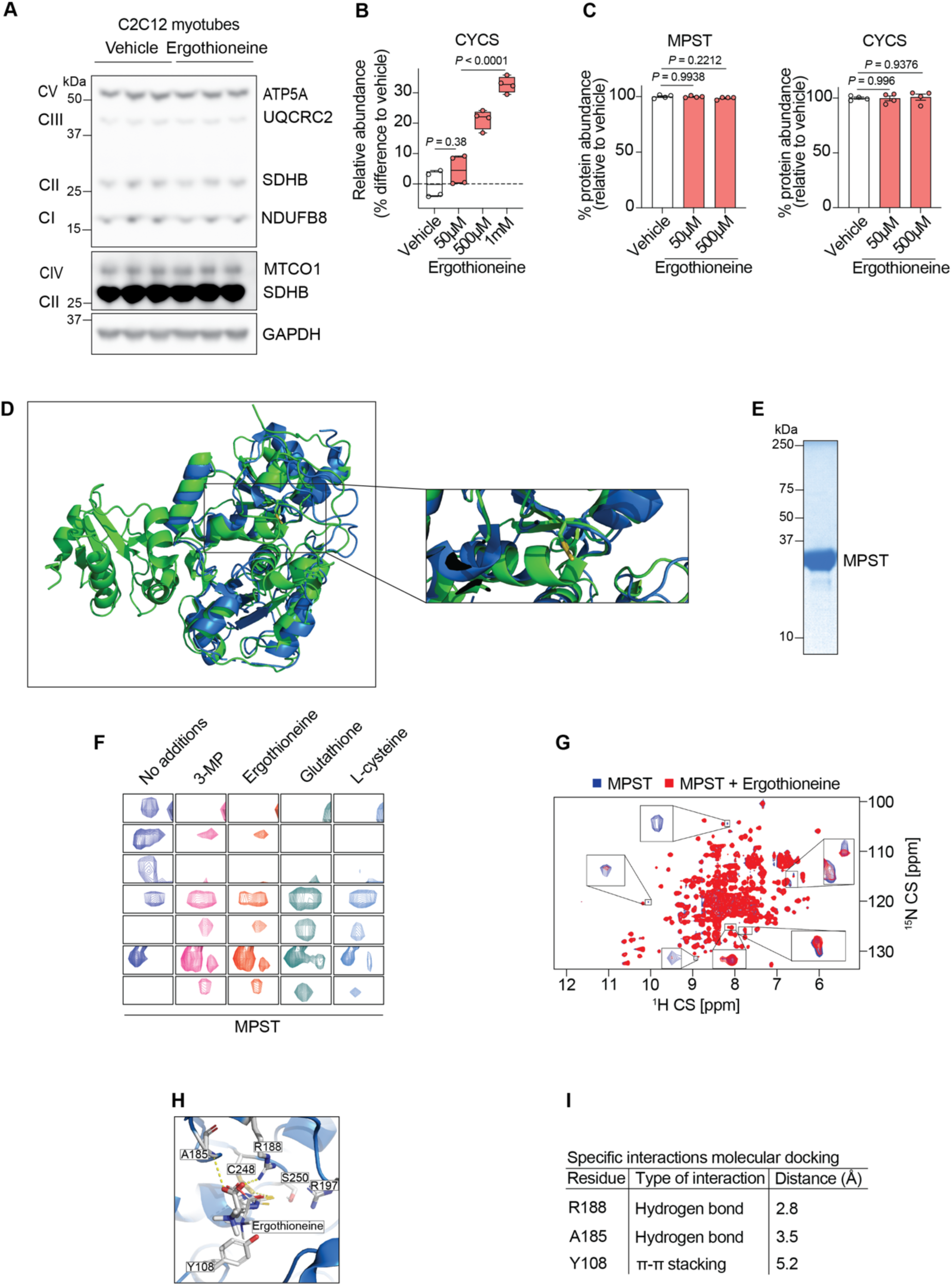
Ergothioneine and MPST. (**A**) Immunoblot analysis of C2C12 myotubes treated with vehicle or 500 µM ergothioneine for 72 h. (**B**) Quantification for CYCS detected in PISA assay across different ergothioneine concentrations (vehicle, 50 µM, 500 µM and 1 mM ergothioneine, n = 4). Center lines denote medians; box limits denote 25^th^ and 75^th^ percentiles; whiskers denote maxima and minima. (**C**) Quantification MPST and CYCS protein levels in whole cell lysates treated with vehicle (n = 4) or different concentrations of ergothioneine (n = 4) overnight. Data are means ± s.e.m.. (**D**) Structural alignment of bacterial ergothioneine synthase Eanb (green, PDB: 6KU1) and human MPST (blue, PDB: 3OLH). (**E**) SDS-gel of ^15^N-labelled recombinant human MPST. (**F**) ^1^H^15^N-HSQC NMR spectra of 50 µM recombinant human MPST (10:1 compound:MPST). (**G**) ^1^H^15^N-HSQC NMR spectra of 600 µM recombinant human MPST (2:1 ergothioneine:MPST). (**H**) Molecular model of the favorable binding mode of ergothioneine in the active site of MPST (PDB: 4JGT). Overlay of predicted poses of ergothioneine are from Glide and nine variants of AutoDock. (**I**) Specific interactions of MPST and ergothioneine predicted by molecular docking analysis. *P* values calculated using one-way analysis of variance (ANOVA).

**Fig. S4.**
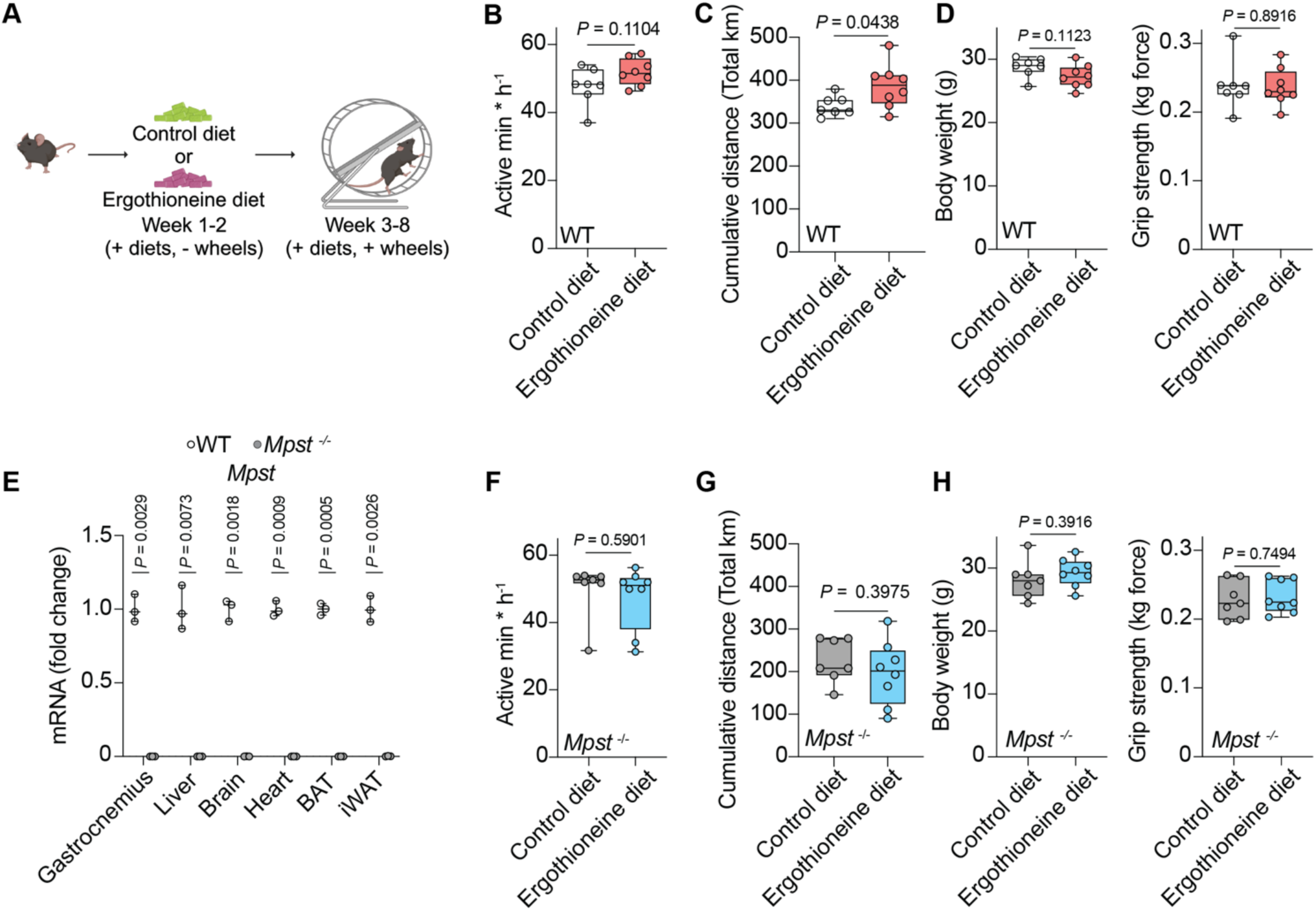
Ergothioneine-enriched diet and exercise training performance. (**A**) Schematic of experimental set up for exercise training experiments upon feeding of control or ergothioneine-enriched diet. Sedentary mice were single housed and fed a control or ergothioneine-enriched diet for 2 weeks. Afterwards exercise training performance was measured using the voluntary wheel running (VWR) system. Mice were exercised for an additional 6 weeks while remaining on the respective diets. (**B**) Active minutes during hour of running speed assessment of control-fed (n = 7) and ergothioneine-fed (n = 8) mice. Data are average values from 3 timepoints. (**C**) Cumulative running distance (km) after week 8 of control-fed (n = 7) and ergothioneine-fed (n = 8) mice. (**D**) Body weight (left panel) and grip strength (right panel) of control-fed (n = 7) and ergothioneine-fed (n = 8) mice after week 8 of the VWR exercise protocol. (**E**) *Mpst* expression in tissues from WT and *Mpst*^-/-^ mice monitored by RT-qPCR (n = 3). (**F**) Active minutes during hour of running speed assessment of *Mpst*^-/-^ control-fed (n = 7) and ergothioneine-fed (n = 8) mice. Data are average values from 3 timepoints. (**G**) Cumulative running distance (km) after week 8 of *Mpst*^-/-^ control-fed (n = 7) and ergothioneine-fed (n = 8) mice. (**H**) Body weight (left panel) and grip strength (right panel) of *Mpst*^-/-^ control-fed (n = 7) and ergothioneine-fed (n = 8) mice after week 8 of the VWR exercise protocol. Center lines denote medians; box limits denote 25^th^ and 75^th^ percentiles; whiskers denote maxima and minima. *P* values calculated using two-tailed unpaired t-test.

